# Coordinated RNA- and protein-templated synthesis of double-stranded DNA by a dual reverse transcriptase immune system

**DOI:** 10.64898/2026.05.04.722794

**Authors:** Megan Wang, Kanta Yoneyama, Rimantė Žedaveinytė, Junichiro Ishikawa, Stephen Tang, Hoang C. Le, Tanner Wiegand, Josephine L. Ramirez, Naoto Nagahata, Yanzhe Ma, Dennis J. Zhang, Erick Helmeczi, Mirela Berisa, Marko Jovanovic, Masahiro Hiraizumi, Keitaro Yamashita, Hiroshi Nishimasu, Samuel H. Sternberg

## Abstract

Defense-associated reverse transcriptase (DRT) systems mediate antiviral immunity through distinct modes of cDNA synthesis: Class 1 DRTs catalyze untemplated synthesis, whereas Class 2 DRTs polymerize noncoding RNA-templated products. However, how these distinct modes drive defense remains unclear. Here, we report that DRT3 immunity arises when Class 1 and Class 2 RT activities cooperate to produce self-complementary double-stranded DNA (dsDNA). DRT3a uses a 5′-ACACAC-3′ RNA template to synthesize poly-(dTdG) repeats, whereas DRT3b synthesizes poly-(dCdA) repeats without any nucleic acid template. Cryo-electron microscopy reveals that DRT3b forms a hexamer and uses active-site-adjacent residues as deoxyadenosine and deoxycytidine gates to enforce alternating nucleotide addition, representing a unique example of amino acid-templated DNA polymerization. DRT3 is toxic in cells lacking RecBCD, implicating host recombination machinery in limiting dsDNA accumulation, and the phage-encoded RecBCD inhibitor Gam triggers DRT3-mediated abortive infection. These findings reveal how two polymerases with distinct templating strategies generate complementary DNA for defense.

## INTRODUCTION

Reverse transcriptases (RTs) are pervasive across bacteria and archaea, where they have been repeatedly co-opted from mobile genetic elements to support diverse pathways of DNA synthesis and host–virus interactions^1^. Phylogenetic analyses indicate that most prokaryotic RTs derive from a limited number of ancestral lineages stemming from group II introns, such as diversity-generating retroelements (DGRs)^2^, and retrons^3^, alongside smaller but rapidly expanding clades such as CRISPR-associated RTs^4^, group II intron-like systems^5,6^, and the so-called Unknown Group (UG) RTs^3^. These lineages illustrate a continuum of functional innovation, from autonomous retroelements capable of mobility, to systems that diversify protein function or generate specialized DNA products, to RNA-templated pathways that interface directly with cellular defense. Notably, many RTs, particularly within UG and abortive infection (Abi) families, function as components of antiviral systems, embedded within modular architectures that couple nucleic acid synthesis to downstream effector activities^1,3^. Together, these observations position RTs as versatile enzymatic platforms for generating diverse DNA products, yet the mechanisms by which these products are formed and translated into cellular outcomes — especially in antiviral immunity — remain incompletely understood.

Unknown Group (UG) reverse transcriptases were initially described as a diverse and poorly characterized lineage^1,3^, but recent genomic and functional studies have established them, alongside independently identified Abi systems, as a major family of defense-associated RTs (DRTs) that often cluster within bacterial defense islands and provide broad protection against bacteriophages^7–14^. Comprehensive phylogenetic analyses have organized UG RT diversity into three major classes distinguished by domain architecture and associated factors^1,3^. Class 1 systems typically encode RTs fused to a characteristic α-helix HEAT-like repeats (αRep) domain, and the systems studied thus far carry out nucleic acid template-independent DNA synthesis, in some cases via protein-primed nucleotide polymerization, as exemplified by AbiK^15^. In contrast, the Class 2 systems investigated thus far rely on conserved non-coding RNAs encoded *in cis* that template the synthesis of defined, tandem-repeat complementary DNA (cDNA) products with diverse downstream functions. Class 3 systems, although less well characterized, are frequently associated with predicted enzymatic effectors such as phosphohydrolases, suggesting additional layers of biochemical coupling between DNA synthesis and effector activity. Across these classes, several members have been experimentally validated as antiphage systems, yet how these distinct modes of DNA synthesis are mechanistically specified and translated into antiviral outcomes remains incompletely understood.

Intriguingly, one UG system — DRT3 — precludes simple classification, as it encodes two reverse transcriptases, DRT3a and DRT3b, that phylogenetically cluster with Class 2 and Class 1 systems, respectively (**Figure 1A**)^3,16^. This unusual genetic architecture raises the possibility that DRT3 integrates fundamentally distinct modes of DNA synthesis within a single immune system, combining RNA-templated and template-independent activities. While recently explored DRT7 systems incorporate an additional DNA polymerizing domain within a single fusion RT polypeptide^17–19^, a biochemical pathway involving two distinct RT enzymes has not yet been described. We therefore set out to investigate how these activities are coordinated, and how their DNA products are integrated to drive antiviral defense.

**Figure 1.**
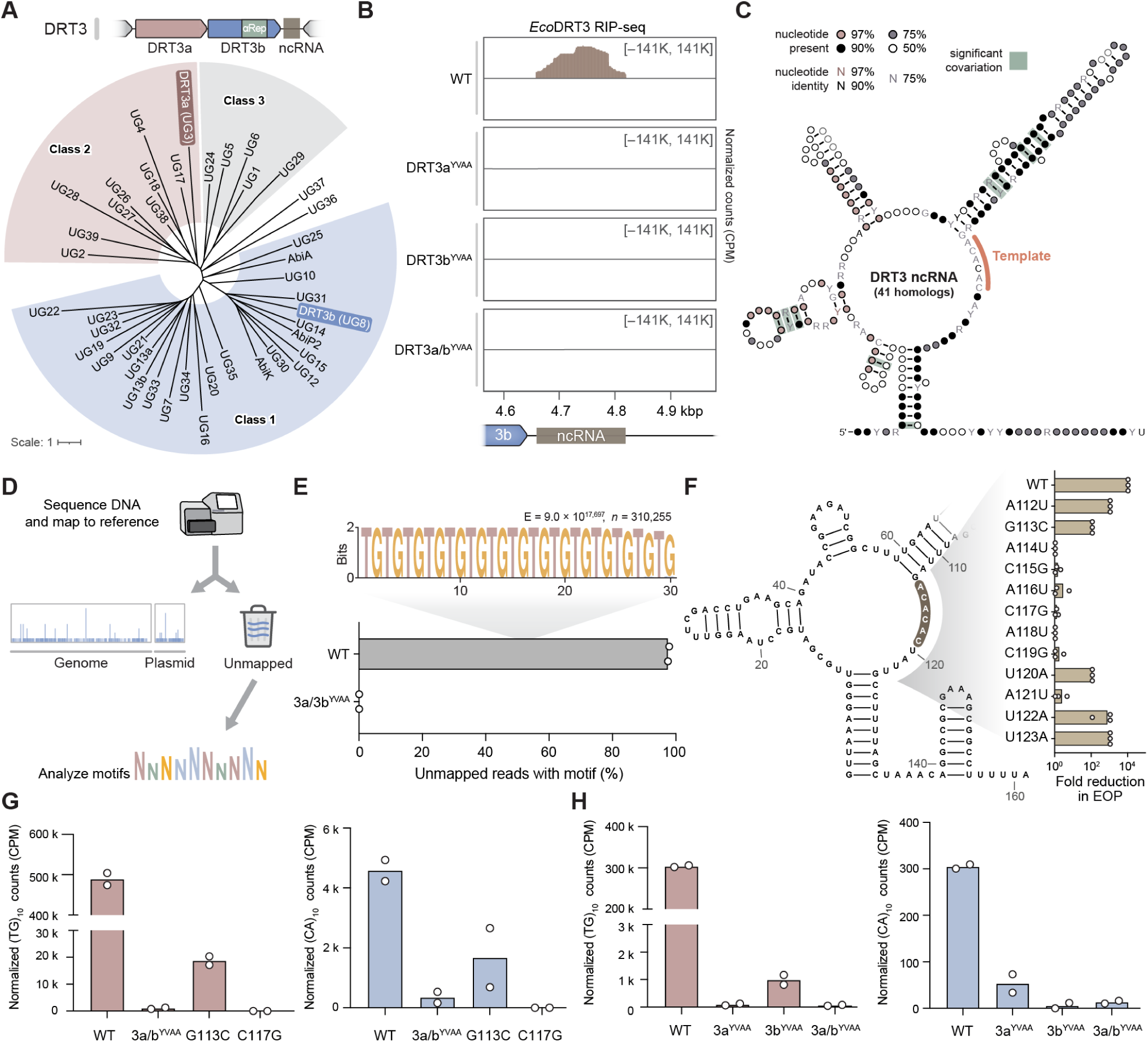
Discovery of RNA-templated DNA synthesis products in the DRT3 system. **(A)** Phylogenetic analysis (bottom) of Unknown Group reverse transcriptases and their clustering into three major clades (Class 1–3), based on a previous study^3^. DRT3 immune systems (top) encode representative RTs from both classes — DRT3a (UG3 within Class 2) and DRT3b (UG8 within Class 1) — as well as a downstream ncRNA. **(B)** RIP-seq profiles across the *Eco*DRT3 locus in WT and catalytic mutant strains using 3×FLAG-tagged DRT3b, showing enrichment of the ncRNA only in the WT background. Coordinates of the ncRNA are displayed below the x-axis. **(C)** Covariance model of the DRT3 ncRNA from an analysis of 41 homologous systems, and WebLogo from a multiple sequence alignment centered around the putative 5′-ACACAC-3′ template region (right). **(D)** Schematic of the analytical pipeline for identifying DNA synthesis products, including sequencing, mapping to reference genomes, and analysis of unmapped reads for enriched sequence motifs. **(E)** Motif analysis of unmapped reads reveals enrichment of poly-(dTdG) repeats in WT samples, which is abolished in catalytic double mutant. **(F)** Functional analysis of ncRNA mutants targeting the conserved 5′-ACACAC-3′ motif-containing loop infected by λ phage, quantified as the fold reduction in efficiency of plating (EOP) relative to an empty vector (EV) control. Data are from n = 3 technical replicates. **(G)** Quantification of normalized cDIP-seq reads containing (dTdG)_10_ (left) and (dCdA)_10_ (right) motifs, plotted as counts per million (CPM), in WT and mutant backgrounds, demonstrating loss of poly-(dTdG) products upon disruption of the ncRNA template. **(H)** Quantification of (dTdG)_10_ and (dCdA)_10_ reads in WT and mutant backgrounds, plotted as in **G**. For **G** and **H**, data represent the mean with individual data points from n = 2 independent biological replicates. See also Figure S1.

Here, we show that DRT3 systems coordinate the synthesis of self-complementary polynucleotides that assemble into double-stranded DNA. DRT3a uses a conserved ncRNA template to generate tandem-repeat poly-(dTdG) species, whereas DRT3b produces complementary poly-(dCdA) strands through a unique protein-templating mechanism that we discovered through cryo-EM analysis. These activities converge to generate double-stranded DNA products whose accumulation is restrained by the host RecBCD pathway, with the phage protein Gam triggering DRT3 immunity by inhibiting RecBCD to enable the buildup of these DNA species and induction of abortive infection. Together, these findings establish DRT3 as a dual-RT system that couples distinct DNA synthesis activities to the production of a double-stranded DNA effector.

## RESULTS

### Discovery of DRT3 cellular DNA synthesis products

We set out to identify DNA reverse transcription products of the DRT3 system in both uninfected and infected cells, building on a cDNA immunoprecipitation and sequencing (cDIP-seq) pipeline previously developed for DRT2, DRT9, and DRT10^7,9,11^. After first recapitulating potent defense against T5 phage for the WT *Eco*DRT3 system heterologously expressed in *E. coli* K-12, as previously described^16^, we appended terminal 3×FLAG tags to either DRT3a or DRT3b, but found that defense activity was completely abolished (**Figure S1A**). To identify permissive tagging sites, we generated AlphaFold3 predictions of each RT enzyme and selected poorly conserved loop regions for internal 3×FLAG insertion, ultimately identifying a site within DRT3b that retained WT phage defense activity (**Figure S1B**). Using this functional tagged variant, we confirmed that defense required intact reverse transcriptase active sites in both DRT3a and DRT3b for protection against T5 and λ phage (**Figure S1C**), and proceeded to test whether the two enzymes physically interact by co-immunoprecipitation followed by mass spectrometry. These experiments revealed a significant interaction between DRT3a and DRT3b, but only when both active sites were intact (**Figure S1D**). Sequencing of their associated RNAs further identified the ncRNA encoded *in cis* downstream of the DRT3b gene, but interestingly, this ncRNA was detected in the immunoprecipitated fraction only when both enzymes were catalytically active (**Figure 1B,C**). Given that DRT3a clusters with Class 2 UG RTs (**Figure 1A**) and is therefore likely to engage the ncRNA directly, these findings suggest that the DNA synthesis products of DRT3a and DRT3b promote formation of a co-complex between both RT enzymes.

We next analyzed DNA species co-purifying with the immunoprecipitated fraction. Conventional read mapping to the ncRNA locus failed to reveal discrete cDNA species (**Figure S1E**), however, based on our previous studies of DRT9 and DRT10, we hypothesized that relevant products would instead be enriched within the unmapped read fraction (**Figure 1D**). Motif analysis of these unmapped reads using MEME identified a prevalent poly-(dTdG) repeat sequence that was uniquely present in cells expressing WT DRT3a and DRT3b (**Figure 1E**). Inspection of the associated ncRNA revealed a highly conserved 5′-ACACAC-3′ sequence within a loop region positioned analogously to the template motifs of DRT9 and DRT10 ncRNAs (**Figure 1C**)^3,9–11^. Consistent with the hypothesis that this region templates poly-(dTdG) synthesis, mutations within this ncRNA segment abolished phage defense and resulted in a complete loss of poly-(dTdG) cDNA products (**Figure 1F,G**). Together, these findings demonstrate that DRT3 mediates RNA-templated synthesis of tandem-repeat cDNAs, similar to other Class 2 UG systems, and suggest that the Class 2 DRT3a enzyme is responsible for this activity.

Our previous work on DRT9 revealed RNA-templated synthesis of poly-dA products, along with complementary poly-dT species that were likely generated by a cellular DNA polymerase^9^. This observation prompted us to more broadly examine the cDIP-seq datasets for additional repeat species beyond poly-(dTdG). Intriguingly, poly-(dCdA) repeats were robustly detected in cells expressing WT DRT3, but not when both RT active sites were mutated (DRT3a/b^YVAA^) (**Figure 1G,H**). Subsequent genetic perturbation experiments revealed a clear division of labor between the two enzymes: poly-(dTdG) repeats were completely abolished when DRT3a, but not DRT3b, was mutated, whereas poly-(dCdA) repeats were specifically lost when DRT3b, but not DRT3a, was mutated (**Figure 1H**). Notably, dinucleotide repeat products were reduced in the presence of either individual RT mutant, suggesting a synergistic effect, while mutation of both active sites eliminated both species (**Figure 1H**). Finally, the levels of both dinucleotide repeats were altered during infection (**Figure S1F**), indicating that phage exposure dynamically influences the activation and/or stability of DNA synthesis products.

Collectively, these results characterized the DNA products generated by DRT3 immune systems, and showed another example of Class 2 UG RTs performing tandem-repeat cDNA synthesis using a confined template portion within associated ncRNA substrates.

### RNA-templated and protein-templated DNA synthesis

To explore the DNA synthesis properties of DRT3 immune systems in greater detail, we recombinantly overexpressed and purified DRT3a and DRT3b separately (**Figure S2**) and assayed their biochemical activities. DRT3a produced RNase-resistant nucleic acid products in the presence of dGTP and dTTP, as assessed by urea–PAGE, and these products were absent when the ncRNA was omitted from the reaction (**Figure 2A**). In contrast, DRT3b selectively generated nucleic acid products in the presence of dATP and dCTP, and this activity was independent of the ncRNA (**Figure 2B**).

**Figure 2.**
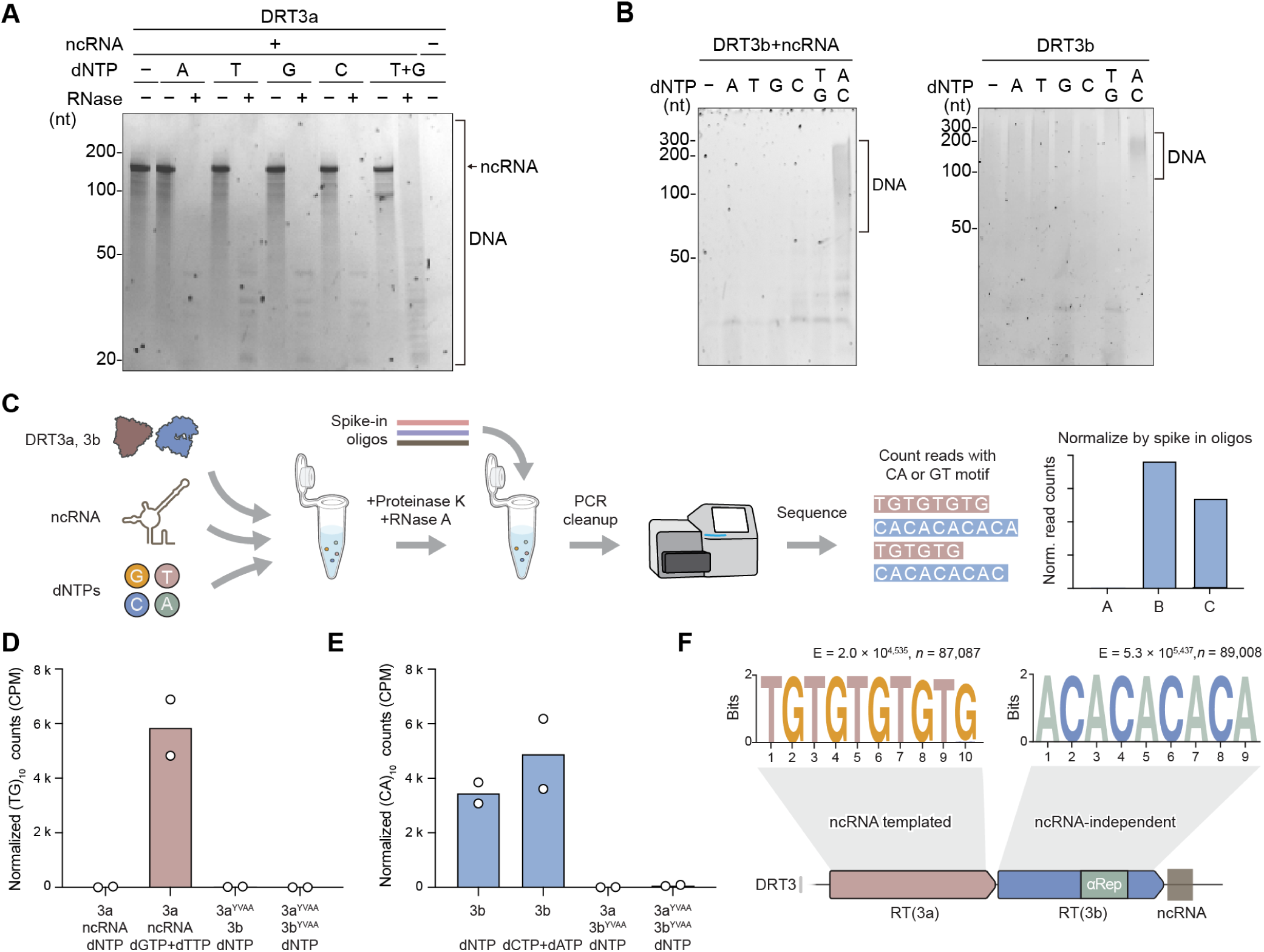
RNA-templated and protein-templated DNA synthesis by DRT3a and DRT3b. **(A)** Urea–PAGE with SYBR Gold stain analysis of *in vitro* DRT3a reactions in the presence of the indicated dNTP combinations with and without the ncRNA. RNase-resistant DNA products are observed specifically in reactions containing dGTP and dTTP and require the ncRNA, consistent with RNA-templated synthesis. **(B)** Urea–PAGE with SYBR Gold stain analysis of *in vitro* DRT3b reactions revealing production of nucleic acid species selectively in the presence of dCTP and dATP. Product formation occurred independent of ncRNA, indicating RNA-independent DNA synthesis. **(C)** Schematic of the DNA high-throughput sequencing (HTS) workflow used to profile *in vitro* DNA synthesis products. Reactions are treated with Proteinase K and RNase A, supplemented with spike-in oligonucleotides, and subjected to HTS and motif-based quantification of poly-(dTdG) and poly-(dCdA) repeat products. **(D)** Sequencing-based quantification of DRT3a-derived products, plotted as normalized (dTdG)_10_ counts per million (CPM). Efficient poly-(dTdG) synthesis is observed only in the presence of ncRNA and dGTP+dTTP and is abolished in catalytic mutant (YVAA) conditions. **(E)** Sequencing-based quantification of DRT3a- and DRT3b-derived products, plotted as normalized (dTdG)_10_ or (dCdA)_10_ counts per million (CPM). Poly-(dCdA) synthesis occurs in the presence of dCTP+dATP or all four dNTPs and is lost upon mutation of RT active sites, whereas productive poly-(dTdG) synthesis requires only dGTP+dTTP. For **D,E**, data represent the mean with individual data points from n = 2 replicate reverse transcription reactions. **(F)** Schematic model summarizing coordinated RNA-templated (DRT3a) and protein-templated (DRT3b) DNA synthesis by the DRT3 system, alongside corresponding Weblogos for both product types derived from the data in **D,E**. Statistical significance of motif enrichment is indicated (E-value), with total reads analyzed (n) shown. See also Figures S2 and S3.

To more sensitively profile DNA products, we developed an alternative experimental approach *in vitro*, where we performed the reverse transcriptase reaction with the purified components of the DRT3 system. After treating them with Proteinase K and RNase A to degrade protein and RNA components, we supplemented the reactions with a defined cocktail of spike-in oligonucleotides, and prepared for subsequent DNA high-throughput sequencing (HTS) and analysis (**Figure 2C**). This approach strongly corroborated our biochemical and cellular findings, confirming that the DRT3a–ncRNA co-complex mediates RNA-templated synthesis of poly-(dTdG) products, while further revealing that the presence of all four dNTP substrates abolished efficient product formation by DRT3a in the absence of DRT3b (**Figure 2D**). In contrast, DRT3b synthesized poly-(dCdA) products independently of ncRNA and was insensitive to the inclusion of all four dNTPs as substrates (**Figure 2E**). Importantly, both product types were completely absent upon mutation of either RT active site (**Figure 2D,E**).

When we analyzed the HTS data more closely, we found that DRT3b products were indistinguishable in length regardless of the nucleotide pool (**Figure S3A**). In contrast, DRT3a–ncRNA products were significantly shorter in the presence of all four dNTPs compared to reactions containing only dGTP and dTTP (**Figure S3A**). Based on the essential role of the ncRNA and its 5′-ACACAC-3′ motif in poly-(dT-dG) synthesis, together with the phylogenetic relationship between DRT3a and DRT10 enzymes^3,11^, we hypothesized that aberrant reverse transcription into stem–loop 4 might drive premature termination of dinucleotide products by disrupting the microhomology required for template resetting (**Figure S3B**). Consistent with this model, motif analysis of HTS read termini revealed an enrichment of sequences corresponding to bases within the stem–loop adjacent to the template region, supporting the idea that readthrough into this structured element results in the observed truncation of the cDNA products (**Figure S3B,C**).

Together, these results establish that DRT3a functions as an RNA-templated reverse transcriptase and produces tandem-repeat poly-(dTdG) cDNA products, whereas DRT3b performs RNA-independent, protein-templated synthesis of poly-(dCdA) DNA products (**Figure 2F**).

### Structural basis for protein-templated poly-(dCdA) synthesis

To elucidate the mechanism of poly-(dCdA) synthesis by DRT3b, we determined its structure by cryogenic electron microscopy (cryo-EM). Preliminary micrographs revealed an oligomeric architecture with apparent threefold symmetry (**Figure S4A**), and subsequent data processing and refinement yielded a 3.1-Å resolution structure of a DRT3b homohexamer (**Figure 3A–D, S4A–D, Video S1**). DRT3b comprises an RT-like domain (residues 1–322), an αRep domain (residues 323–621), and a C-terminal tail (CTT; residues 622–667) (**Figure 3A–D, Video S2**). Unlike canonical reverse transcriptases^20^, the RT-like domain contains palm and finger subdomains but lacks an α-helical thumb. Instead, the N-terminal region of the αRep domain occupies the position typically filled by the thumb domain. The αRep domain adopts an α-solenoid fold composed of six repeating α-helical units and mediates oligomerization, while the CTT forms a loop at the interface between the RT-like and αRep domains (**Figure 3D**). Interactions between the αRep domain of one protomer and the RT-like domain of an adjacent protomer promote trimer formation, with two trimers assembling into a hexamer primarily through RT–RT interactions (**Figure 3C**). This oligomeric architecture resembles other hexameric bacterial RT assemblies, including AbiK^15^, DRT4^21^, and DRT9^9,10^, which likewise form higher-order complexes through protein–protein interfaces, although in contrast to DRT9 — where the ncRNA plays a central scaffolding role — DRT3b appears to assemble independently of nucleic acid cofactors.

**Figure 3.**
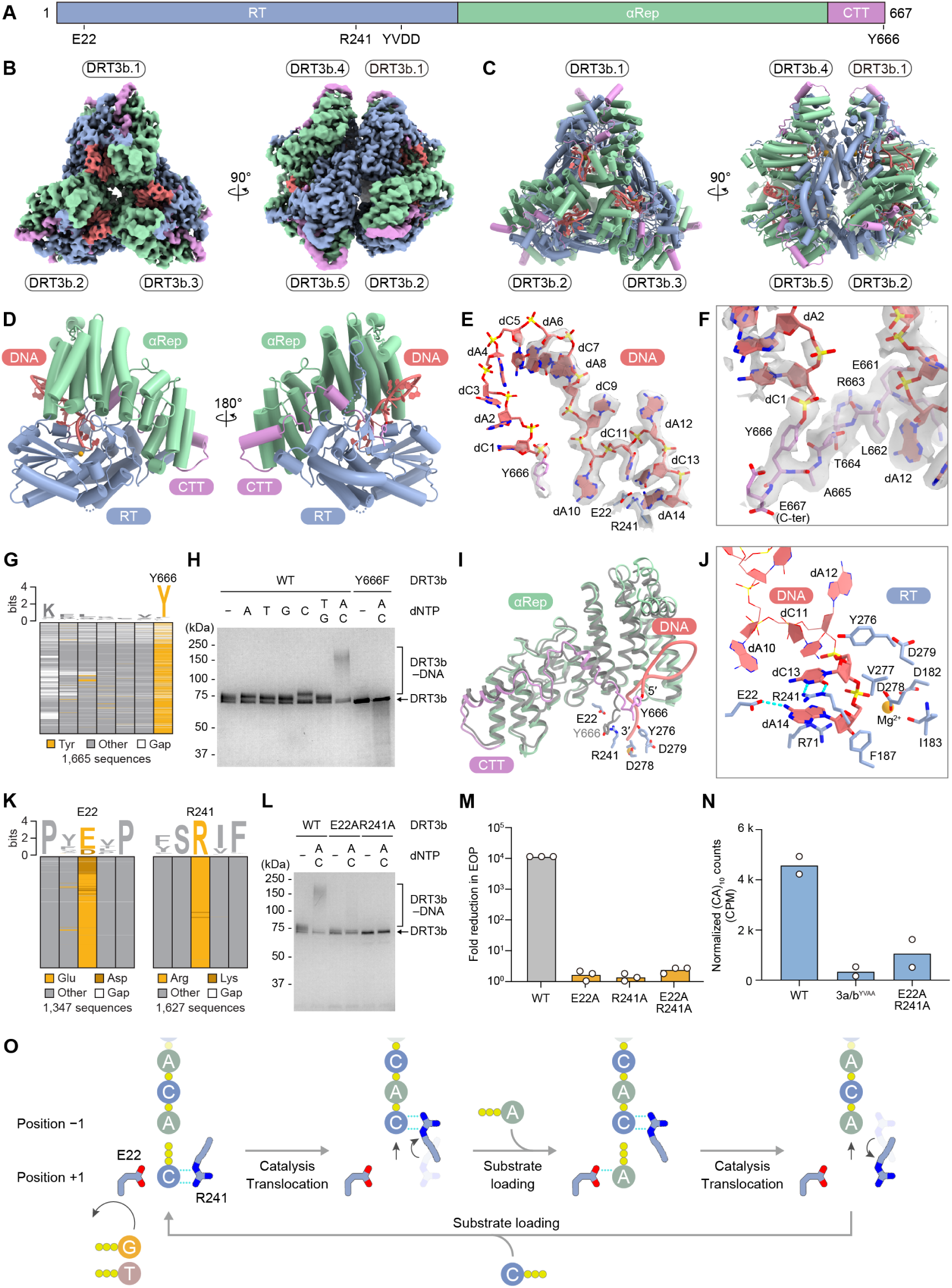
Structural basis of DRT3b-catalyzed poly-(dCdA) synthesis. **(A)** Domain architecture of DRT3b, showing the RT-like domain, αRep domain, and C-terminal tail (CTT). Conserved residues implicated in DNA synthesis and nucleotide selectivity are indicated. **(B)** Cryo-EM density of the DRT3b hexamer, shown in two orientations. Domains are colored as in **A**, and DNA density is shown in red. Protomers are labeled to highlight the dimer-of-trimers architecture. **(C)** Cartoon representation of the DRT3b hexamer in the same orientations and colors as in **B**. **(D)** Cartoon representation of a single DRT3b protomer in two orientations, showing a bound DNA synthesis product within the catalytic cleft. **(E)** Structure of a 14-mer (dCdA)_7_ strand. Individual nucleotides are labeled, with the 5′ end extending toward the side chain of Y666. Cryo-EM densities are shown as a semi-transparent gray surface for the DNA and key residues (E22, R241, and Y666). Nucleotides dC1–dC5 are less well resolved in the density map, likely due to their flexibility. **(F)** Close-up view of the DNA 5′ end extending to the side chain of Y666, with surrounding C-terminal tail residues shown. Cryo-EM densities are displayed at lower contour level to highlight the Y666–dC1 linkage. **(G)** Sequence conservation of the DRT3b C-terminal region across homologs, highlighting the invariant Y666 residue. **(H)** SDS–PAGE analysis of DRT3b DNA synthesis reactions with the indicated dNTPs. A higher-mobility species corresponding to a covalent DRT3b–DNA adduct is observed with the WT enzyme in the presence of dCTP and dATP, but is abolished in the Y666F mutant. **(I)** Comparison between the DNA-bound DRT3b structure and an AlphaFold3 model of DRT3b in the DNA-free state (colored gray), indicating that Y666 is positioned at the active site for priming DNA synthesis. Key residues in or near the active site are labeled. **(J)** Close-up view of the DRT3b active site showing key interactions between the DNA product and residues E22 and R241, as well as nearby residues in the active site. Hydrogen bonds are depicted with dashed cyan lines. **(K)** Sequence conservation of E22 (left) and R241 (right) across DRT3b homologs. **(L)** SDS–PAGE analysis of DRT3b DNA synthesis reactions in the presence of dATP and dCTP, showing a loss of the covalent DRT3b–DNA adduct in the presence of E22A or R241A mutations. **(M)** Functional analysis of protein templating residues, quantified as the fold reduction in efficiency of plating (EOP) relative to an EV control. Data are from n = 3 technical replicates. **(N)** Quantification of (dCdA)_10_ reads in WT and mutant backgrounds, plotted as counts per million (CPM). Data represent the mean of n = 2 independent biological replicates. **(O)** Mechanistic model of DRT3b-catalyzed poly-(dCdA) synthesis. E22 functions as a gatekeeper to exclude dTTP and dGTP and facilitate dATP incorporation, while R241 serves as a specificity switch to facilitate dCTP incorporation through conformational changes, thereby enforcing alternating nucleotide addition. Hydrogen bonds are depicted with dotted cyan lines. See also Figures S3-S7, Videos S1-S3, and Table S5.

Closer inspection of the cryo-EM map revealed conspicuous densities that fit well to a polynucleotide with an alternating purine–pyrimidine sequence (**Figure 3E, S4E**). Notably, this density was continuous from the 5′ end to the side chain of Y666 (**Figure 3F**), a residue that is highly conserved at the primary sequence level (**Figure 3G**), suggesting that, similar to other Class 1 UG RTs^15,22^, the DNA product is protein-primed and covalently attached to DRT3b via this residue. Consistent with this interpretation, SDS–PAGE analysis of DRT3b in the presence of dNTP substrates revealed a higher-mobility species when dCTP and dATP were included, and this species was abolished by a Y666F mutation that removes the hydroxyl group required for priming (**Figure 3H**). cDIP–seq of DRT3-containing cells further showed that poly-(dCdA) products — but not poly-(dTdG) products — were completely eliminated in the presence of the DRT3b^Y666F^ mutant, and defense against λ phage was likewise abolished (**Figure S5A,B**). Finally, DNA synthesis reactions performed with [α–^32^P]dATP produced a radioactive species that migrated at the expected molecular weight of the DRT3b protein by denaturing SDS–PAGE (**Figure S5C**), indicating a covalent protein–DNA adduct. Together, these findings demonstrate that DRT3b functions as a protein-primed reverse transcriptase that directly extends DNA from the side chain of Y666. Interestingly, AlphaFold3 modeling of the DNA-free state positioned Y666 immediately adjacent to the DRT3b active site, consistent with a pre-catalytic configuration for protein-primed DNA synthesis (**Figure 3I**).

We modeled the polynucleotide density as a 14-mer stretch of (dCdA)_7_ (**Figure 3E**), consistent with our *in vitro* and *in vivo* DNA sequencing analyses. Notably, the samples used for cryo-EM structure determination were not incubated with dNTP substrates, indicating that DNA products were generated during protein overexpression and likely represent co-purifying ligands resistant to degradation by cellular nucleases. To gain mechanistic insight into nucleotide selectivity, we examined conserved residues surrounding the active site and identified R241 and E22 — both highly conserved across diverse DRT3b homologs — as candidate determinants of alternating dCTP and dATP incorporation, respectively (**Figure 3J,K, S5D, Video S3**). Mutational analysis supported this model, as individual substitutions in either residue resulted in a loss of signal for DNA-linked protein gel-shift and phage defense (**Figure 3L,M**), while combined mutations abolished both defense and detectable increases in poly-(dCdA) synthesis from the catalytic mutant (**Figure 3M,N**).

Structural analysis of the DRT3b active site further revealed that dA14 and dC13 occupy the binding pockets for the incoming nucleotide (position +1) and the 3′ end of an elongating DNA strand (position −1), respectively (**Figure 3J, S4F**). The dA14 nucleobase engages in hydrogen-bonding and stacking interactions with E22 and R71, respectively, while its ribose moiety contacts F187 (**Figure 3J, S5D**). Additionally, its phosphate group is stabilized by a Mg^2+^ ion coordinated by D182, I183, and D278 within the YVDD motif. In turn, the dC13 nucleobase forms bidentate hydrogen bonds with R241, with its ribose moiety interacting with Y276 and V277 (**Figure 3J, S5D**). Modeling alternative nucleotides at position +1 revealed electrostatic repulsion between the carbonyl groups of thymine or guanine and the negatively charged side chain of E22 (**Figure S6A**), explaining the inability of DRT3b to incorporate dTTP or dGTP. Unlike adenine, the modeled cytidine at position +1 is too distant (3.9 Å) from E22 to form a hydrogen bond (**Figure S6A**), explaining why dATP, but not dCTP, is preferentially incorporated at position +1 when cytidine occupies position −1 and interacts with R241. Furthermore, R241 would sterically clash with the modeled adenine at position −1, thereby moving from position −1 toward position +1 (**Figure S6B**). Consequently, dCTP, but not dATP, is selectively accommodated at position +1 when adenine occupies position −1 (**Figure S6B**).

Like the E22A and R241A mutants, the E22K and R241E mutants exhibited almost no activity *in vitro*, further confirming the essential roles of E22 and R241 in DNA synthesis (**Figure S6C**–**S6E**). To assess whether substitutions in these critical residues remain catalytically active *in vivo*, we sequenced the DNA products recovered from *E. coli* cells expressing DRT3 carrying E22 and R241 substitutions that occur in other DRT3 homologs (**Figure 3K**). Notably, the E22D variant retained partial poly-(dCdA) synthesis, consistent with this substitution occurring in a substantial fraction of DRT3b homologs, whereas R241K, present in a non-significant subset of homologs, produced no detectable product (**Figure S6F**). Meanwhile, the R71A and R71K mutations reduced DNA synthesis, suggesting that R71 contributes to defining the nucleotide preference (**Figure S6G**). Although both R71A and R71K retained partial synthesis *in vitro*, only R71K produced the detectable (CA)_10_ motif in cells (**Figure S6H**). This result likely reflects that R71A’s residual activity is too low to leave detectable products in cells, whereas R71K, which retains a long aliphatic side chain, remains more active than R71A *in vivo*.

Together, these results suggest that (1) E22 and R241 facilitate the incorporation of dATP and dCTP, respectively, through hydrogen-bonding interactions, (2) E22 excludes dTTP and dGTP while permitting dCTP, and (3) R241 shifts its position between positions −1 and +1, thereby enforcing alternating nucleotide incorporation. This framework also explains the lower-efficiency synthesis of poly-dC relative to poly-(dCdA) (**Figure 3H**). Collectively, these findings demonstrate that DRT3b catalyzes amino acid–dependent poly-(dCdA) synthesis through coordinated actions of E22 as a gatekeeper and R241 as a specificity switch (**Figure 3O**).

Finally, we identified residues that mediate key interactions within the trimeric DRT3b interfaces as well as the dimer-of-trimers interface (**Figure S7A,B**). Mutations disrupting the trimer interface resulted in a complete loss of phage defense, whereas perturbations at the dimer-of-trimers interface had minimal effects, retaining near-WT activity (**Figure S7C**). Consistent with these observations, direct quantification of DNA synthesis products revealed that trimer interface mutations largely eliminated both poly-(dTdG) and poly-(dCdA) species (**Figure S7D**). Together, these results establish that DRT3b functions as an oligomeric, protein-templated reverse transcriptase in which conserved amino acid side chains both prime DNA synthesis and enforce the production of alternating adenosine–cytidine dinucleotide repeats.

### Synergistic DRT3a and DRT3b activities produce dsDNA

We next sought to explore the coordinated synthesis of poly-(dTdG) and poly-(dCdA) by DRT3a–ncRNA and DRT3b complexes, respectively. We were particularly interested in whether these self-complementary polynucleotides could form double-stranded DNA, and whether this behavior might explain — or arise from — higher-order interactions between both modules, consistent with their co-precipitation in RIP-seq and proteomics analyses (**Figure 1B, S1D**).

We began by monitoring DNA synthesis product yields in biochemically reconstituted reactions containing DRT3a–ncRNA, DRT3b, or both. As observed before (**Figure S3A**), normalized levels of po-ly-(dTdG) products were low when DRT3a–ncRNA was incubated with all four dNTPs, but increased dramatically upon addition of DRT3b — regardless of whether DRT3b harbored an intact active site (**Figure 4A**). In contrast, poly-(dCdA) product levels were largely insensitive to the presence of DRT3a–ncRNA (**Figure 4A**). Analysis of read length distributions further revealed that poly-(dCdA) species were substantially longer in the presence of DRT3a–ncRNA, independent of its catalytic activity, whereas poly-(dTdG) products synthesized by DRT3a approached their maximum length only when catalytically active DRT3b was present (**Figure 4B**). Together, these results indicate that DRT3a and DRT3b function synergistically, with each enzyme enhancing the synthesis and/or elongation of the other’s preferred DNA products, consistent with coordinated production of complementary strands.

**Figure 4.**
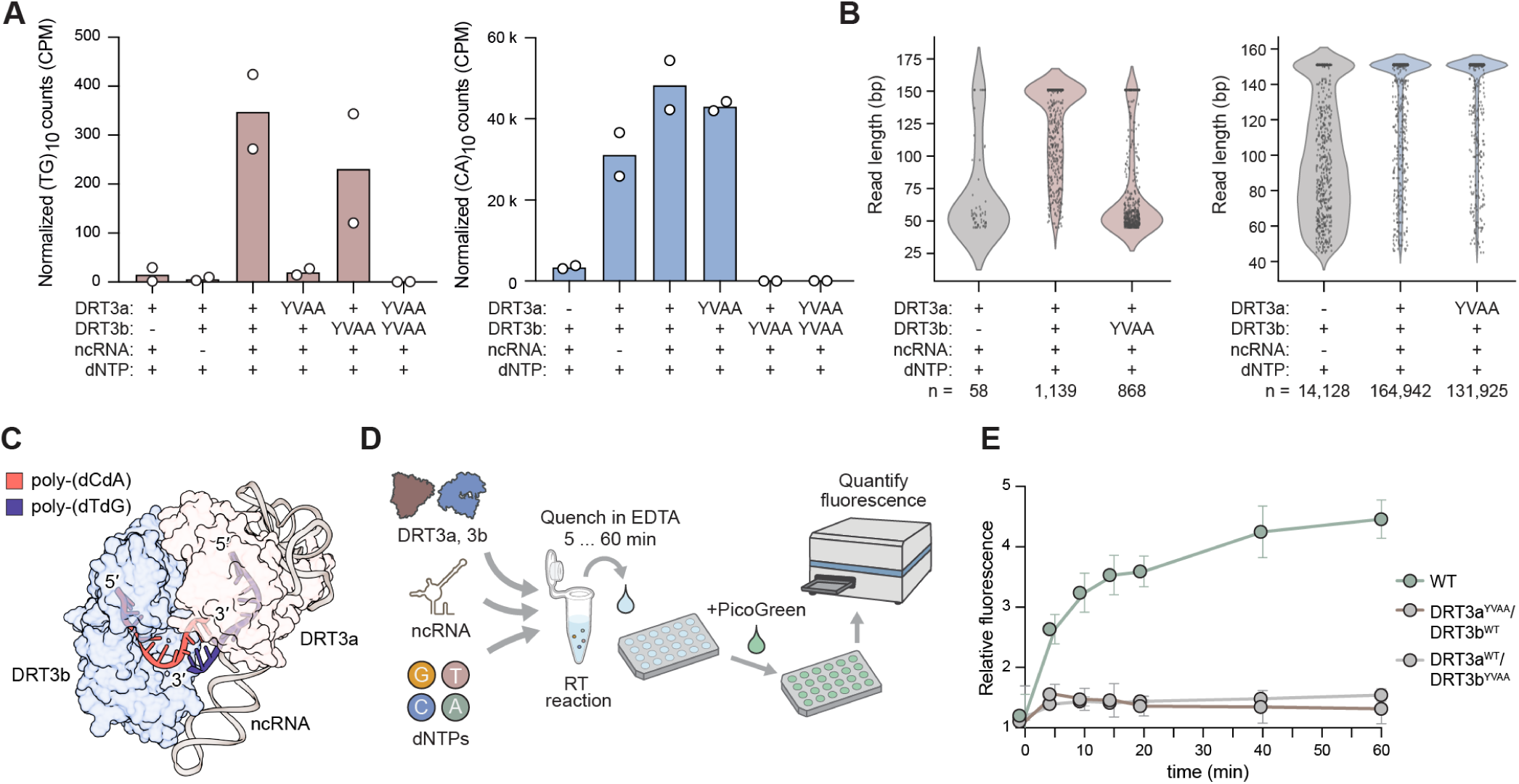
Coordinated DRT3a and DRT3b activities drive complementary DNA synthesis and dsDNA formation. **(A)** Quantification of DRT3a- and DRT3b-derived products, plotted as normalized (dTdG)_10_ or (dCdA)_10_ counts per million (CPM). Poly-(dTdG) synthesis by DRT3a is strongly enhanced by the presence of DRT3b, including a catalytically inactive mutant, whereas poly-(dCdA) synthesis by DRT3b is largely unaffected by DRT3a. Data represent the mean with individual data points from n = 2 replicate reverse transcription reactions. **(B)** Read length distributions of DNA products generated under the indicated conditions. Left, DRT3a-derived products; right, DRT3b-derived products. DRT3b products are longer in the presence of DRT3a–ncRNA independent of its catalytic activity, whereas DRT3a products reach maximal length only when catalytically active DRT3b is present. The numbers of reads (n) are indicated below each condition and up to 500 individual data points were plotted per condition. **(C)** Alphafold3 model illustrating coordinated synthesis of complementary poly-(dTdG) and poly-(dCdA) products by DRT3a–ncRNA and DRT3b, respectively, and their potential annealing into double-stranded DNA. **(D)** Schematic of the PicoGreen-based assay used to monitor dsDNA formation during *in vitro* reverse transcription. Reactions are quenched at the indicated time points, incubated with PicoGreen, and fluorescence is measured to quantify double-stranded DNA. **(E)** PicoGreen fluorescence measurements showing time-dependent accumulation of dsDNA in reactions containing WT DRT3a and DRT3b, but not in reactions containing catalytic mutants (YVAA). Fluorescence is plotted relative to reactions containing DRT3a and DRT3b catalytic mutants at the corresponding time point. Data are mean ± SD of n = 3 separate reverse transcription reactions. See also Figure S8.

Building on these findings, we next tested whether DRT3 generates double-stranded DNA through the combined activity of DRT3a–ncRNA and DRT3b (**Figure 4C**). Using a PicoGreen-based assay to monitor dsDNA formation — which leverages the strong fluorescence increase of PicoGreen upon binding double-stranded, but not single-stranded, nucleic acids (**Figure 4D**) — we observed a robust, time-de-pendent signal increase in reactions containing catalytically active DRT3a and DRT3b, even in the absence of dedicated hybridization steps, consistent with progressive dsDNA accumulation (**Figure 4E**). In contrast, reactions containing individual catalytic mutants failed to produce signal above background (**Figure 4E**). These results provide direct evidence that coordinated DRT3a–DRT3b activity drives the formation of complementary DNA products that anneal into dsDNA, supporting a model in which dsDNA accumulation underlies DRT3-mediated defense.

Of note, our biochemical analyses failed to identify a specific priming residue for the DRT3a-me-diated DNA synthesis products, in contrast to the protein-primed mechanism we identified for DRT3b. Denaturing SDS–PAGE and urea–PAGE analyses did not reveal higher-mobility protein or ncRNA species under conditions that promoted poly-(dTdG) synthesis (**Figure S8A,B**), nor did incubation with [α–^32^P] dGTP yield a detectable radioactive DRT3a band greater than the size of the ncRNA (**Figure S8C**). We also failed to observe DNA-primed DNA synthesis using short oligonucleotide primers, and re-analysis of HTS data failed to reveal any evidence of poly-(dTdG) products extending directly off of DRT3b-me-diated poly-(dCdA) products (**Figure S8D,E**). These results suggest that DRT3a initiates DNA synthesis through an alternative mechanism, potentially involving *de novo* initiation with a dNTP substrate.

### Phage escapers implicate RecBCD in DRT3 regulation

To gain further insight into the immune function of double-stranded poly-(dCdA)/(dTdG) polymers and the mechanism of DRT3 activation, we sought to isolate phage mutants that escape DRT3 immunity. Using high-titer plaque assays, we identified rare escape plaques for λ phage, amplified them in a WT DRT3 strain, and retested their sensitivity to DRT3 defense (**Figure 5A**). From these experiments, we isolated nine λ mutants that robustly escaped DRT3 defense (**Figure S9A**), and whole-genome sequencing revealed a diverse set of loss-of-function mutations that all mapped to a single gene: *gam* (**Figure 5B**). Consistent with this result, recombinant expression of WT *gam* was sufficient to activate DRT3 immunity, inducing rapid and pronounced growth arrest when co-expressed with DRT3, but not in a mutant DRT3 or mutant *gam* background (**Figure 5C, S9B–D**). This phenotype mirrored the growth arrest observed during phage infection at high multiplicity of infection (**Figure 5D**), supporting a model in which DRT3 immunity induces abortive infection and programmed dormancy.

**Figure 5.**
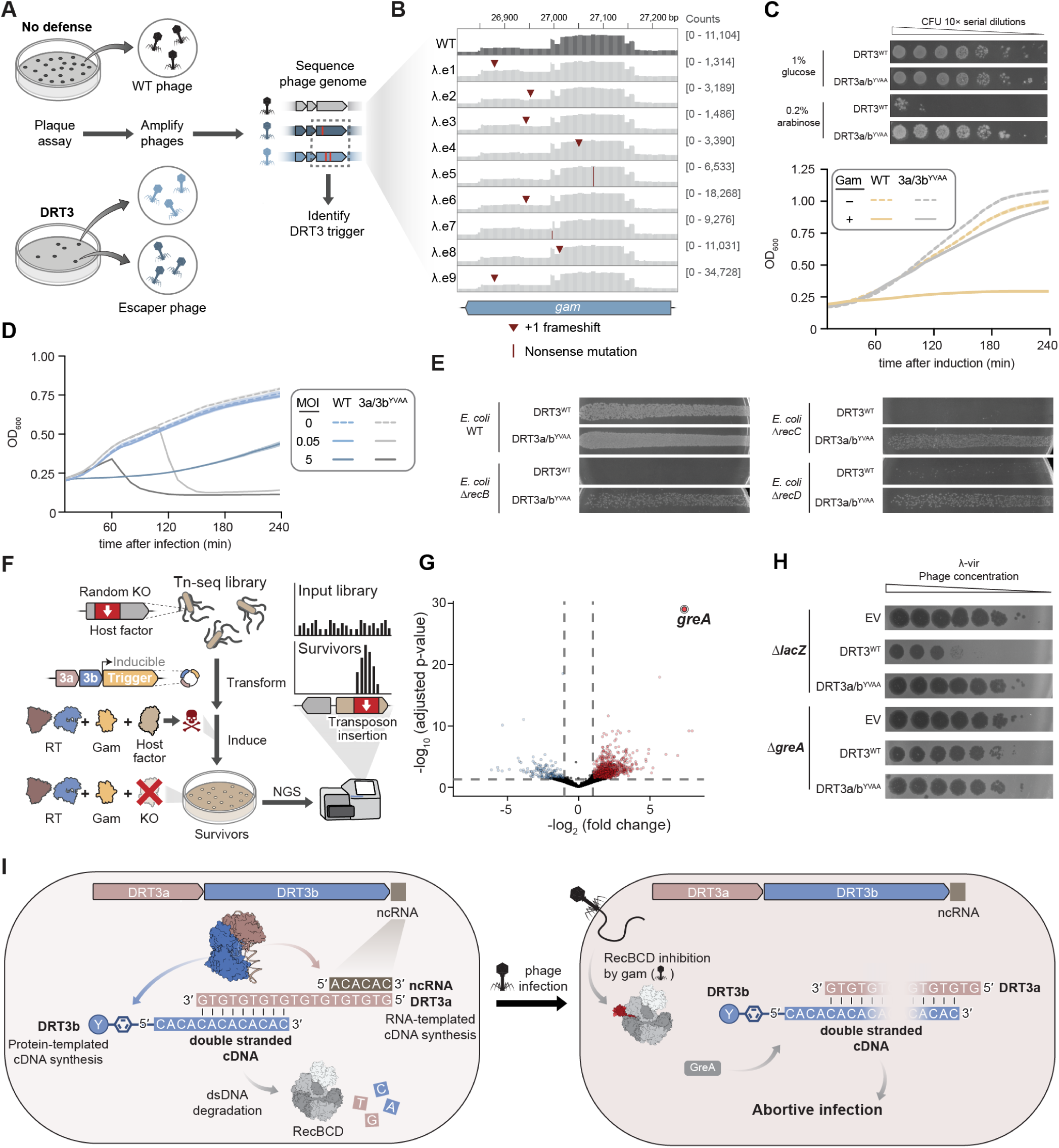
RecBCD inhibition by λ phage Gam activates DRT3-mediated defense. **(A)** Schematic of the genetic selection used to identify phage triggers of DRT3. Cells lacking DRT3 (no defense) or expressing DRT3 were infected with phage and plated to isolate escaper mutants. Escapers were amplified and their genomes sequenced to identify mutations associated with evasion of DRT3-mediated defense. **(B)** Mutational profiling of independent phage escapers mapped to the λ *gam* locus. Read coverage across the genomic region is shown for WT and individual escaper isolates, with mutations indicated. Recurrent +1 frameshift and nonsense mutations in *gam* were identified across independent isolates. **(C)** Cell viability and growth assays showing that induction of *gam* is toxic in the presence of DRT3. Top, cell viability assay under inducing conditions. Bottom, growth curves of cells expressing wild-type or mutant *gam* from an inducible expression vector, monitored over time after induction of *gam* or an empty vector control. Shaded regions indicate the SD across independent biological replicates (n = 3). **(D)** Growth curves following λ phage infection at the indicated multiplicities of infection (MOI) in cells expressing DRT3. Phage infection results in growth inhibition in a DRT3-dependent manner, consistent with an abortive infection immune response. Shaded regions indicate the SD across independent biological replicates (n = 3). **(E)** Plaque assays comparing phage replication on cells expressing WT or mutant DRT3 in the indicated genetic backgrounds. Loss of RecBCD function causes synthetic lethality in DRT3-expressing strains. **(F)** Schematic of the transposon mutagenesis screen followed by high-throughput sequencing (Tn–seq), to identify host factors required for DRT3- and Gam-mediated toxicity. After generating a high-density transposon mutant library, cells were transformed with a plasmid encoding DRT3 and Gam, and then subjected to selection under inducing conditions. Surviving clones were analyzed by high-throughput sequencing to identify enriched transposon insertions. **(G)** Volcano plot of Tn–seq results from **F** showing differential enrichment of transposon insertions, under DRT3 + Gam induction conditions relative to DRT3 catalytic mutant + Gam induction. Each point represents a gene, and *greA* is among the most significantly enriched hits. Dashed lines indicate thresholds for statistical significance (p_adj_ < 0.05) and fold-change cutoffs (|log_2_FC| > 1). **(H)** Plaque assays assessing λ-vir phage replication on cells expressing empty vector, WT DRT3, or a catalytic mutant (DRT3a/ b^YVAA^) in a Δ*greA* knockout background or a control Δ*lacZ* background. Loss of *greA* results in a loss of DRT3-mediated defense. **(I)** Model for DRT3 activation by phage-encoded triggers and regulation by RecBCD. Under uninfected conditions, RecBCD degrades DRT3-derived DNA products, preventing their accumulation. Upon phage infection, inhibition of RecBCD by Gam allows these DNA products to accumulate, promoting DRT3-mediated abortive infection and restriction of phage replication. See also Figures S9-S11, Tables S6 and S7.

Phage λ Gam is a well-characterized inhibitor of the host RecBCD complex (also known as ExoV), a multifunctional exonuclease–helicase that processes dsDNA ends during homologous recombination, while also serving as a key antiviral factor that degrades incoming phage dsDNA during infection. This connection prompted us to test whether DRT3 activity depends on RecBCD function. Strikingly, we were unable to propagate the WT DRT3 expression plasmid in *E. coli* strains containing genetic knockouts in any of the RecBCD components (**Figure 5E**). These findings strongly implicate RecBCD as a regulatory factor that enables persistent and constitutive DRT3 activity, likely by recognizing and degrading its double-stranded poly-(dCdA)/(dTdG) products. This model parallels our recent finding that ExoI (the *sbcB* gene product), a host ssDNA-specific exonuclease, prevents premature accumulation of DRT9-derived poly-dA species^9^. Consistent with a shared principle, side-by-side experiments revealed selective synthetic lethality between DRT3 and loss of RecBCD (Δ*recB*), and between DRT9 and loss of ExoI (Δ*sbcB*), but not alternative combinations (**Figure S9E**). These results highlight a unifying role for host nucleases in controlling the accumulation and activity of RT-derived DNA synthesis products.

Next, we wanted to explore the possible functions of poly-(dCdA)/(dTdG) products and the mechanism of abortive infection. We first tested the hypothesis that unchecked DNA synthesis might deplete intracellular nucleotide pools required for both cellular growth and phage genome replication. However, metabolomic profiling failed to detect any depletion of key adenosine- or thymidine-derived nucleotides in either uninfected or infected cells, indicating that DRT3 activity does not function through nucleotide exhaustion (**Figure S10A**). Reasoning that transcriptomics might reveal pathways activated during DRT3 immunity, we performed RNA–seq on λ phage-infected cells with or without a functioning DRT3 system and identified a large set of differentially enriched genes, including a number of up-regulated genes within the SOS response pathway^23^, such as *recA*, *lexA*, and *sulA* (**Figure S10B, S10D**). However, an *E. coli* strain lacking RecA — a central initiator of the SOS response — retained robust DRT3-mediated defense (**Figure S10C**), suggesting that this pathway is not essential for the abortive infection phenotype.

Reasoning that an unbiased approach would be more effective in identifying as yet unknown pathways that interface with DRT3-mediated abortive infection, we performed a transposon mutagenesis screen to identify host factors required for DRT3 function^24^. In brief, we used the *Mariner* transposon to construct a high-saturation mutagenesis library in *E. coli*, transformed with a single-plasmid vector encoding DRT3 and Gam. We then selected surviving clones under induction conditions that cause cell death in a WT background, and resulting populations were analyzed using high-throughput sequencing (**Figure 5F**). Consistent with our prior observations, comparing DRT3 to DRT3a/b^YVAA^ under Gam suppression revealed significant depletion of transposon insertions in only a single gene, *recC* (**Figure S10E**). Negative controls encoding catalytically inactive DRT3a/b compared to DRT3 wild-type system, revealed strong enrichment of transposon insertions in a small set of loci, with *greA* emerging as the most significant hit (**Figure 5G**). GreA is a conserved transcription factor that rescues backtracked RNA polymerase and promotes transcriptional fidelity, and its absence has been implicated in enhanced DNA break repair via RecBCD-mediated resection^25^. To validate the involvement of GreA in DRT3-mediated immunity, we performed phage defense assays in a Δ*greA* background and found that loss of GreA led to a potent loss of DRT3-mediated defense (**Figure 5H**), without any detectable effects on levels of DRT3 expression (**Figure S10F**).

Collectively, our results provide multiple lines of evidence that the RecBCD recombination machinery regulates DRT3-based dsDNA products, and further implicate transcription-associated processes in DRT3-mediated abortive infection.

## DISCUSSION

A central paradigm in molecular biology is that organisms with double-stranded DNA genomes propagate their genetic material via semiconservative replication, in which each parental strand serves as a template for the synthesis of its complementary strand by DNA polymerase. Here, we show that the DRT3 immune system challenges this paradigm by generating dsDNA through two mechanistically distinct and independently templated processes. The DRT3a reverse transcriptase uses a noncoding RNA to produce repetitive poly-(dTdG) cDNA products, analogous to bacterial DRT10 and eukaryotic telomerase machineries^11^, whereas the DRT3b reverse transcriptase employs a protein-primed and protein-templated mechanism to produce precise poly-(dCdA) dinucleotide repeats. Despite these fundamentally distinct templating strategies and modes of substrate engagement — with DRT3a accommodating an RNA–DNA hybrid and DRT3b acting on single-stranded DNA — both enzymes operate using the same underlying reverse transcriptase fold and catalytic chemistry. Remarkably, their self-complementary products spontaneously anneal to form double-stranded DNA, whose accumulation is required to drive abortive infection (**Figure 5I**).

While this system represents a striking departure from canonical DNA replication, the sequence of the dsDNA product itself is intriguing. The alternating purine-pyrimidine repeats within the poly-(dCdA)/ (dTdG) duplex under specific conditions form Z-DNA^26,27^, an atypical left-handed double helix that adopts a more rigid and constrained conformation than canonical, right-handed B-DNA^28^. In bacteria, Z-DNA formation has been linked to genetic instability and spontaneous deletion events that revert the causative sequences to a state more compatible with B-form DNA, presumably reflecting the cellular cost of stably maintaining the left-handed conformation^29,30^. Intriguingly, Z-form nucleic acids function as an immunity sensor in mammalian cells via the factor Z-DNA binding protein 1 (ZBP1), which senses the structured nucleic acid and triggers necroptosis, apoptosis, and interferon signaling^31,32^. Whether DRT3-derived DNA products adopt a Z-DNA conformation *in vivo* — and whether this feature contributes to, or causes, abortive infection — will be important to resolve.

Consistent with the idea that the dsDNA product of DRT3 functions as an immune signal in the cell, our results revealed an active role for host pathways in constraining its accumulation. We identified the phage λ Gam protein as a trigger of DRT3-mediated toxicity, and in turn, established that the host RecBCD complex functions as a key suppressor of this activity. Genetic inactivation of any RecBCD subunit phenocopied Gam expression, leading to DRT3-dependent cell death. In our model, RecBCD continuously degrades DRT3-synthesized DNA prior to infection, keeping levels below the threshold necessary to cause cellular dormancy. Upon infection, phage-encoded DNA mimic proteins like Gam inhibit RecBCD to alleviate this suppression and activate DRT3-mediated abortive infection, mirroring the ‘guard’ functions of RecBCD in other diverse defense systems^18,33,34^. Notably, this requirement is strand-selective: loss of dsDNA exonuclease activity was synthetically lethal with DRT3, whereas disruption of the ssDNA-specific exonuclease SbcB had no comparable effect. Together, these findings establish DRT3 as a unique defense system in which accumulation of a dsDNA product, rather than its synthesis alone, activates abortive infection.

To probe the underlying mechanism, we took an unbiased genetics approach, reasoning that host factors required for DRT3-mediated toxicity might provide key insights. Through a transposon mutagenesis screen, we unexpectedly identified GreA as a host factor required for DRT3-mediated defense. GreA is a transcription factor that canonically rescues backtracked RNA polymerase to promote transcriptional fidelity^35^, a function not readily connected to DRT3-mediated DNA synthesis. However, this observation becomes striking in light of recent work from Gao and colleagues that identified the phage T1 protein ST61 as a trigger of DRT3 activation^36^. Although its function remains unknown, ST61 is predicted to adopt an SH3-like fold that has predicted structural similarity to the transcription factors GreA and Spt4/5 (**Figure S11**), which are known to interact with host RNAP. Together, these observations raise the exciting possibility that DRT3 activation is linked to perturbations of the cellular transcription machinery, potentially through interference of RNAP function.

The most surprising finding of our study lies in the enzymatic mechanism of DNA synthesis itself, a result that corroborates elegant complementary work from Deng et al.^36^. Our structural and biochemical analyses establish that DRT3b initiates DNA synthesis through a protein-primed mechanism, in which the conserved Y666 residue is covalently linked to the nascent DNA product. The cryo-EM structure further revealed an unexpected mode of protein-templated poly-(dCdA) synthesis, in which active site residues enforce selective incorporation of strictly alternating dCTP and dATP nucleotides. Although protein-directed nucleotide selection has been observed in a limited number of systems — most notably in the trans-lesion polymerase Rev1, tRNA CCA-adding enzymes, and terminal deoxynucleotidyl transferase (TDT) enzymes^37–39^ — these mechanisms operate in fundamentally different contexts and lack the precise repeating periodicity of DRT3b. Our observations raise the possibility that similarly unconventional modes of DNA synthesis may be widespread among Class 1 UG RT systems, many of which remain functionally uncharacterized, with unknown DNA products and sequence specificities.

Alongside recent work, our findings point to a broader landscape in which reverse transcriptases encode diverse, exotic, and unanticipated mechanisms of DNA synthesis. DRT3 highlights a previously unappreciated capacity for these enzymes to encode protein-directed rules of DNA synthesis, and to couple these outputs to cellular fate. The continued discovery of unexpected activities at the host–virus interface underscores how existing molecular frameworks can be creatively repurposed through evolutionary exaptation to produce unanticipated biological functions. These observations suggest that similarly unconventional chemistries — and their functional consequences — remain hidden across microbial genomes, awaiting discovery.

## LIMITATIONS OF THE STUDY

While our study defines the molecular logic of DRT3-mediated antiphage defense, including the ncRNA- and amino acid-templated synthesis of its unusual DNA products, the terminal effector mechanism by which DRT3 activation leads to abortive infection remains incomplete. Our results establish that DNA synthesis from both RTs is required for defense, and our genetic data exclude an SOS–RecA–SulA axis, but we cannot yet assign what leads to growth arrest. We found that *greA* deletion abolished defense, thus implicating this transcription elongation factor, but further experiments will be needed to determine whether it functions upstream of DRT3, as part of the activating signal, or downstream of DNA synthesis. Furthermore, the synergistic effects between DRT3a and DRT3b require further exploration. Our sequencing analyses together with co-immunoprecipitation results indicate that both enzymes act synergistically to generate their respective DNA products. However, the molecular basis of this cooperation, and how the activities of the two enzymes are coordinated, remains to be defined. Finally, it will be important to identify the specific priming mechanism employed by the DRT3a enzyme.

## Supporting information

Key Resources Table

## RESOURCE AVAILABILITY

### Lead contact

Further information and requests for resources and reagents should be directed to the lead contact, Samuel H. Sternberg.

### Materials availability

The study did not generate any new materials or reagents.

### Data and code availability

Next-generation sequencing data will be made available in the National Center for Biotechnology Information (NCBI) Sequence Read Archive (BioProject accession: PRJNA1461384) and the Gene Expression Omnibus (GSE330639, GSE330641, GSE329894, GSE329895, GSE329897) at the time of publication. Atomic coordinates have been deposited in the Protein Data Bank under the accession code 26CZ. The cryo-EM density map has been deposited in the Electron Microscopy Data Bank under the accession code EMD-80550. The raw images have been deposited in the Electron Microscopy Public Image Archive under the accession code EMPIAR-13689. IP–MS data are available in the MassIVE database under accession number MSV000101674. Other datasets generated and analyzed in the current study are available from the corresponding authors on reasonable request. Custom scripts used for bioinformatics, RNA–, RIP–, cDIP–, miniprep–seq, *in vitro* reverse transcription sequencing, and Tn–seq data analyses are available upon request.

## ACKNOWLEDGMENTS

We thank A.J. Robinson, A.I. Palmieri, R. Rafat, and T.M. Smith for laboratory support; the JP Sulzberger Columbia Genome Center for NGS support; staff scientists at The University of Tokyo’s cryo-EM facility, especially Yoichi Sakamaki, for help with cryo-EM data collection; and Dr. Makoto Nakakido and Dr. Kouhei Tsumoto for mass photometry analysis; A. Thawani for guidance on radioactive assays; E. Richard, G.D. Lampe, and all the members of the Sternberg lab for valuable discussions. Metabolomics work was performed at the Metabolomics Core at the Icahn School of Medicine at Mount Sinai (RRID:SCR_027540). M.W. was supported by NIH Medical Scientist Training Program grant (5T32GM145440). S.T. was supported by a Ruth L. Kirschstein Individual Predoctoral Fellowship (F30AI183830) from the NIH. M.J. was supported by NIH grant R35GM152258. H.N. was supported by JSPS KAKENHI Grant Numbers 21H05281 and 25H00436, JST CREST Grant Number JPMJCR23B6, the Takeda Medical Research Foundation, and the Inamori Research Institute for Science. S.H.S. was supported by NSF Faculty Early Career Development Program (CAREER) Award 2239685, a Pew Biomedical Scholarship, an Irma T. Hirschl Career Scientist Award, a Mallinckrodt Scholarship, the Howard Hughes Medical Institute Investigator Program, and a generous startup package from the Columbia University Irving Medical Center Dean’s Office and the Vagelos Precision Medicine Fund.

## AUTHOR CONTRIBUTIONS

S.T., R.Z., M.W., and S.H.S. conceived the project. M.W. performed DRT3 defense assays, cDIP–seq experiments, miniprep–seq experiments, biochemical experiments analyzed by high-throughput sequencing, radioactive assays, dsDNA formation assays, and computational analysis of sequencing assays. K.Yoneyama and J.I. purified recombinant proteins and performed biochemical experiments analyzed electrophoretically. K.Yoneyama, J.I., N.N., M.H., K.Yamashita, and H.N. performed structural analyses. R.Z. performed RNA–seq, cDIP–seq experiments, cell viability, and trigger co-expression assays. S.T. performed RNA–seq, RIP–seq, and cDIP–seq experiments, phage escaper screening and validation experiments, RT phylogenetic analyses, and ncRNA covariance modeling. H.C.L. performed Tn-seq, RNA–seq, and GreA functional assays, with support from M.W. T.W. performed phylogenetic analyses and assisted in data interpretation and visualization. J.L.R. performed early cloning and DRT3 defense experiments, RIP-seq assays, and initial protein expression tests. Y.M. and M.W. performed proteomics experiments, with supervision from M.J. D.J.Z. performed early DRT3 defense experiments and bioinformatics analyses. E.H. and M.W. performed metabolomics experiments and analyses, under the guidance of M.B. M.W., K.Yoneyama, R.Z., H.N., and S.H.S. discussed the data and wrote the manuscript, with input from all authors. H.N. and S.H.S. supervised the project.

## STAR METHODS

### EXPERIMENTAL MODEL AND STUDY PARTICIPANT DETAILS

#### Bacterial strains and phages

*Escherichia coli* strains (**Table S1**) (MG1655, BW25113, DE3, and Δ*sbcB*, Δ*recA,* Δ*recB*, Δ*recC*, Δ*recD,* Δ*sulA*, Δ*greA* from the Keio collection^40^) were grown in LB or LB agar at 37 °C shaking at 300 rpm. Whenever applicable, media were supplemented with chloramphenicol (20 µg mL^−1^) or kanamycin (50 µg mL^−1^) to ensure the maintenance of plasmids (**Table S2**).

The phages used in this study are listed in the **Key Resources Table**. Infection was performed in LB medium or in soft LB agar supplemented with 5 mM CaCl_2_ and 5 mM MgSO_4_ at 37°C as explained in Method Details.

#### Method Details

##### Plasmids and strain construction

The *Eco*DRT3 operon encoding *Eco*DRT3a and *Eco*DRT3b (**Table S3**) with its native upstream and downstream flanking sequences were chemically synthesized (Twist Bioscience) and cloned into the pACYC184 plasmid backbone via Gibson Assembly. Variant plasmids were cloned through a variety of methods, including around-the-horn PCR, restriction digestion–ligation, and Golden Gate assembly. All plasmids were cloned and propagated in *E. coli* strain NEB Turbo (sSL0410), and clones were verified by Sanger or whole plasmid sequencing via Genewiz or Plasmidsaurus. Experiments were performed in *E. coli* K-12 strain MG1655 unless otherwise indicated.

##### Phylogenetic analyses

The phylogenetic tree displaying classes of UG RT homologs in **Figure 1A** was plotted using data from a previous study^3^. Each UG/DRT clade was collapsed into a single representative, using the longest branch within each clade for ease of visualization. To build the phylogenetic tree of DRT3a RT homologs in **Figure S3B**, *Eco*DRT3a was used as the seed for a BLASTp search of the NCBI NR database (E-value threshold 0.05, max target sequences = 5,000). Hits were clustered at 95% sequence identity using MMseqs2^41^ and aligned using MAFFT^45^ v7.490. A phylogenetic tree was constructed from the resulting alignment with FastTree 2^43^ (-wag -gamma) v2.1.11, prior to visualization and annotation in iTOL^44^.

##### ncRNA covariance modeling

A covariance model for the DRT3-associated ncRNA was built as previously described^9^. Putative ncRNAs associated with *Eco*DRT3 and closely related systems (subgroup 9 from Mestre et al.^3^) were analyzed by retrieving 250 nt of intergenic sequence downstream of each operon. Nucleotide sequences were aligned using MAFFT^45^ (v7.490) and trimmed to the boundaries of the *Eco*DRT3 ncRNA identified by RIP-seq. These sequences were realigned using mLocARNA^46^ (v2.0.1) with default parameters. The resulting multiple sequence alignment was used to build and calibrate a covariance model (CM) using Infernal^47^ (v1.1.4). Next, a larger database of DRT3 homologs was generated using *Eco*DRT3b as the seed for a BLASTp search (E-value threshold 0.05, max target sequences = 5,000). Nucleotide sequences corresponding to each hit, along with 1 kb of upstream and downstream sequence, were extracted and clustered at 90% identity using CD-HIT^48^. CMsearch from the Infernal suite was used to interrogate the resulting sequence database to identify additional ncRNA hits and refine the model, yielding 32 unique ncRNA hits. The final CM was evaluated for statistically significant covariation and visualized using R-scape^49^ (v2.6.10) at an E-value threshold of 0.05 in “Evaluate given structure” mode.

##### Phage propagation and plaque assays

Phages T5 and λ-vir were gifts from M. Laub. All phages were propagated in liquid culture by picking single plaques into LB medium containing *E. coli* MG1655 cells diluted 1:100 from overnight cultures, supplemented with 5 mM CaCl_2_ and 5 mM MgSO_4_. After 3–4 hours of shaking incubation at 37 °C, chloroform was added to a final concentration of 5% to lyse any residual cells and then pelleted via centrifugation at 4,000*g* for 5 min. The resulting supernatant was filtered using a sterile 0.22-µm filter and stored at 4 °C.

Small-drop plaque assays were performed as previously described^9^. Briefly, *E. coli* K-12 MG1655 (sSL0810) (strain descriptions and genotypes are provided in **Table S2**) were transformed with the indicated plasmids (plasmid descriptions and sequences are provided in **Table S1**). Transformants were inoculated into a culture of LB medium and antibiotic, with overnight shaking at 37 °C. The next day, overnight cultures were mixed with molten soft agar (0.5% agar in LB medium supplemented with 5 mM CaCl_2_ and 5 mM MgSO_4_), cooled to 45 °C, and then plated on solid-bottom agar (1.5% agar in LB medium containing the appropriate antibiotic) in a Petri dish. Then, 10× serial dilutions of phage in LB medium were spotted onto the surface of the soft agar lawn. The plates were incubated at 37 °C for 8–16 hours to allow for plaque formation. Plaque-forming units (PFU)/mL were calculated using the following formula:. When individual plaques were indistinguishable, a count of 50 plaques was assigned at the lowest dilution with visible bacterial lawn clearance. Phage defense activity was assessed by fold-reduction in efficiency of plating (EOP), which was calculated by dividing the PFU/mL formed on a lawn of the empty vector containing condition by the PFU/mL formed on a lawn of the defense system-containing condition.

##### RIP–seq and cDIP–seq

RIP–seq and cDIP–seq were performed as previously described^9^. *E. coli* K-12 strain MG1655 (sSL0810) cells were transformed with plasmids encoding the DRT3 operon that is either WT, single, or double mutant (YVAA) RT active site with a DRT3b internal 3×FLAG tag (plasmid sequences are provided in **Table S1**). Individual colonies for each replicate experiment were inoculated in 40 mL liquid LB and grown at 37 °C to an OD_600_ of 0.3–0.4. For experiments performed with phage infection, cultures were split in half and either infected with λ-vir or T5 in LB medium at an MOI of 5 or with the same volume of LB for 20 minutes. For experiments performed with uninfected cells, 20 mL of cultures were grown to an OD_600_ of 0.4 and collected. Cells were then immediately pelleted by spinning down at 4,000*g* for 5 minutes at 4 °C, and then washed with 5 mL of cold TBS (20 mM Tris-HCl pH 7.5 at 25 °C, 150 mM NaCl). Cells were then spun down again at 4,000*g* for 5 minutes at 4 °C. The pellets were washed with 1 mL of cold TBS and centrifuged at 10,000*g* for 5 min at 4 °C. After removing supernatants, the pellets were flash-frozen in liquid nitrogen and stored at −80 °C.

To prepare antibody–bead complexes for immunoprecipitation, protein G Dynabeads (Thermo Fisher Scientific) were washed 3 times with 1 mL of IP lysis buffer (20 mM Tris pH 7.5 at 25 °C, 150 mM KCl, 1 mM MgCl_2_, 0.2% Triton X-100). Washed beads were then resuspended in 1 mL of IP lysis buffer and combined with anti-Flag antibody (Sigma-Aldrich, F3165) and rotated at 4 °C for at least 3 hours. 60 µL of beads and 20 µL of antibody mix were prepared for each sample. Then, antibody–bead complexes were washed 3 times in 1 mL of IP lysis buffer to remove unconjugated antibodies and resuspended to a final volume of 60 µL per sample.

Flash frozen pellets were thawed on ice and resuspended in 1.2 mL IP lysis buffer supplemented with 1× Xpert Protease Inhibitor Cocktail Solution (GenDEPOT) and 0.1 U µL^−1^ SUPERase•In RNase Inhibitor (Thermo Fisher Scientific). Cells were lysed by sonication and lysates were cleared by centrifugation at 21,000*g* for 15 min at 4 °C. The supernatants were transferred to separate tubes and two 10 µL aliquots of samples were set aside for inputs (10 µL each for RIP and cDIP) and stored at −80 °C. 60 µL of antibody–bead conjugates were added to each remaining immunoprecipitation sample and rotated overnight at 4 °C. After overnight incubation, the samples were washed 3 times with 1 mL IP wash buffer (20 mM Tris-HCl pH 7.5 at 25 °C, 150 mM KCl, 1 mM MgCl_2_). During the final wash, each sample was split into two 500 µL volumes for downstream RIP or cDIP processing.

For RIP, the supernatants were removed and beads were resuspended in 750 µL TRIzol (Thermo Fisher Scientific), while inputs were resuspended in 740 µL and incubated for 5 min at room temperature. The supernatant containing eluted RNA was taken into new tubes with 150 µL of chloroform and incubated at room temperature for 2 min before centrifugation at 12,000*g* for 15 min at 4 °C. The 400 µL of the aqueous phase was taken into a new tube and then RNA was cleaned up with the RNA Clean & Con-centrator-5 kit (Zymo Research). cDIP elution was performed by removing the supernatant, resuspending beads with 80 µL of IP wash buffer containing 5 µg RNase A (Thermo Fisher Scientific). The inputs were resuspended with 70 µL of the same wash buffer. Both inputs and immunoprecipitation samples were incubated at 37 °C for 30 minutes. Then, SDS was added to a final concentration of 1% along with 25 µg of proteinase K (Thermo Fisher Scientific). The supernatants containing eluted DNA were transferred to new tubes and DNA was isolated with the Monarch Spin PCR and DNA Cleanup kit (NEB), according to the Oligonucleotide Cleanup protocol.

For RIP–seq library preparation, RNA was diluted in NEBuffer 2 and heated at 92 °C for 2 min for fragmentation by alkaline hydrolysis. The samples were then treated with TURBO DNase (Thermo Fisher Scientific), RppH (NEB), and T4 PNK (NEB) to remove DNA and prepare RNA ends for adapter ligation. RNA was purified using the Zymo RNA Clean & Concentrator-5 kit. Adapter ligation, reverse transcription, and indexing PCR were performed using the NEBNext Small RNA Library Prep kit. Libraries were sequenced on an Element AVITI system in single or paired-end mode with 75 or 150 cycles per end.

For cDIP–seq library preparation, the samples were heated at 95 °C for 2 min and then immediately placed on ice to denature DNA. Adapter ligation and conversion of ssDNA to dsDNA were performed using the xGen ssDNA & Low-Input DNA Library Prep Kit (IDT). Indexing PCR was performed using NEBNext Ultra II Q5 Master Mix. Libraries were sequenced on the Element AVITI system in single or paired-end mode with 75 or 150 cycles per end.

##### Miniprep–seq

Miniprep–seq samples were prepared as previously described^11^. In brief, *E. coli* K-12 MG1655 (sSL0810) was transformed with a plasmid encoding the *Eco*DRT3 operon together with λ *gam* under a pBAD-inducible promoter. Overnight cultures grown in 1% glucose were diluted 1:100 in 80 mL LB medium supplemented with 1% glucose and grown until OD_600_ 0.3 before spinning down at 4,000*g* for 5 min and resuspending in 80 mL LB medium. 15 mL of the culture was taken out as time point 0 and the rest of the culture was grown in LB supplemented with 0.2% arabinose. 15 mL of culture was taken out at each time point and immediately spun down at 4,000*g* for 5 min. After removing the supernatant at each time point, cell lysis and DNA extraction was immediately performed following the standard protocol for the QIAprep Spin Miniprep Kit (QIAGEN) and elution in 30 µL nuclease-free water. Sequencing libraries were prepared as described above for cDIP and sequenced on the Element AVITI platform in single- or paired-end mode at 75 or 150 cycles per end.

##### RIP–seq analysis

RIP–seq data were analyzed as previously described^9^. Briefly, raw reads were processed with Cutadapt^50^ (v5.0) to filter out adapter sequences, trim low-quality bases, and remove reads shorter than 15 bp. Filtered reads were aligned with bwa-mem2^51^ (v2.2.1) under default parameters to a composite reference comprising the *E. coli* MG1655 genome (NCBI: NC_000913.3), the relevant plasmid sequence, and either the λ-vir (NCBI: NC_001416.1) or T5 (NCBI: NC_005859.1) phage genome. Alignments were sorted and indexed using SAMtools^52^ (v1.17). Strand-specific coverage tracks were generated with bamCoverage^53^ (v3.5.6), normalized to sequencing depth based on the total number of reads passing trimming and length filtering. Coverage tracks were visualized in IGV^54^.

Reads from total RNA–seq libraries were assigned to annotated transcriptome features with featureCounts^55^ (v.2.0.2), using the -s 1 flag to account for library strandedness. The resulting count matrices were passed to PyDESeq2^56^ (v.0.5.3) with default parameters, which estimated per-transcript fold-change and false-discovery rate between sample conditions indicated in the figure legend. Results were displayed as volcano plots generated in matplotlib, with log_2_[fold change] plotted as a function of −log_10_[p_adj_]. Three independent biological replicates were included for every comparison.

##### Total RNA–seq

*E. coli* K-12 MG1655 was transformed with the plasmids listed in **Table S1** and cultured overnight in LB containing the appropriate antibiotics and 1% glucose. Overnight cultures were back-diluted 1:100 into 7 mL of LB with 1% glucose, grown to OD_600_ 0.25, and pelleted by centrifugation at 4,000*g* for 5 min. Pellets were washed once with 20 mL of LB to remove residual glucose, then resuspended in 7 mL of fresh LB. Each suspension was divided into 3 mL aliquots supplemented with antibiotics and either 1% glucose or 0.2% arabinose. Following shaking incubation at 37 °C for 15 min, the cultures were spun down at 4,000*g* for 5 min and the supernatant was discarded, and the pellet was resuspended in 750 µL TRIzol and incubated for 5 min at room temperature. Next, 150 µL of chloroform was added to each sample, incubated for 2 min at room temperature, and then spun down at 12,000*g* for 15 min at 4 °C. Next, 360 µL of the aqueous phase was transferred to a clean tube and purified with the RNA Clean & Concentrator-5 kit (Zymo Research). Library preparation for total RNA–seq samples was performed as described above for RIP samples.

##### cDIP–seq, *in vitro* biochemical sequencing, miniprep–seq analysis

Sample reads were processed by Cutadapt^50^ (v5.0) to filter out adapter sequences, trim low-quality bases, and remove reads shorter than 15 bp. Specifically, Cutadapt was run using a tiling approach across the 3′ adapter, as previously described^57^, with parameters set to -e 0.2, -O 10, -q 30, and -m 45 (error rate, minimum overlap, quality cutoff, and minimum length, respectively). For cDIP**–**seq and miniprep**–**seq, filtered reads were then aligned with bwa-mem2^51^ (v2.2.1) under default parameters to a composite reference comprising the *E. coli* MG1655 genome (NCBI: NC_000913.3), the relevant plasmid sequence, and either the λ-vir (NCBI: NC_001416.1) or T5 (NCBI: NC_005859.1) phage genome. For *in vitro* biochemistry sequencing experiments, trimmed reads were aligned to a reference containing the sequences of the three 150-bp spike-in oligonucleotides. Alignments were sorted and indexed using SAMtools^52^ (v1.17). Strand-specific coverage tracks were generated with bamCoverage^53^ (v3.5.6), normalized to sequencing depth based on the total number of reads passing trimming and length filtering. Coverage tracks were visualized in IGV^54^.

##### Unmapped read analysis

Reads from cDIP–, miniprep–seq, and *in vitro* biochemical sequencing that failed to align with bwa-mem2^51^ (v2.2.1) were extracted using SAMtools fasta^52^. The fraction of unmapped reads was calculated using SAMtools flagstat. Unmapped reads were analyzed with MEME^58^ (v5.5.7) to identify enriched sequence motifs. Specifically, MEME was run in differential enrichment mode with the any number of repetitions model and a minimum motif width of 6 bp, using unmapped reads from each sample of interest as the positive set and unmapped reads from the catalytically inactive RT (YVAA) control as the negative set. Motif-containing reads were quantified using a custom Python script. Each read was screened for a stretch of ≥10 consecutive TG or CA bases, allowing a Hamming distance of 1. The number of reads containing a hit was then normalized by dividing by the total number of trimmed or unmapped reads for cDIP–seq, plasmid-mapped reads for miniprep–seq, or aligned reads for *in vitro* biochemistry sequencing.

##### Co-IP–MS

For immunoprecipitation, *E. coli* K-12 strain MG1655 cells were transformed with the *Eco*DRT3 plasmid and its double catalytic mutant DRT3a/b^YVAA^ containing an internal 3×FLAG tag on DRT3b. Overnight cultures were diluted 1:100 in 50 mL of LB medium and grown until OD_600_ 0.4. Cells were pelleted by spinning down 4,000*g* for 10 min at 4 °C. The supernatant was removed before washing the pellet with 1 mL cold TBS, spun down again 15,000*g* for 1 min at 4 °C, and pellets were flash-frozen in liquid nitrogen.

To prepare antibody–bead conjugates, protein G Dynabeads (Thermo Fisher Scientific) were washed 3 times with 1 mL of IP–MS lysis buffer (20 mM Tris pH 7.5 at 25 °C, 150 mM KCl, 1 mM MgCl_2_, 0.2% Triton X-100). Washed beads were then resuspended in 1 mL of IP–MS lysis buffer and combined with anti-Flag antibody (Sigma-Aldrich, F3165) and rotated at 4 °C for at least 3 hours. 100 µL of beads and 10 µL of antibody mix were prepared for each sample. Then, antibody–bead complexes were washed 3 times in 1 mL of IP–MS lysis buffer to remove unconjugated antibodies and resuspended to a final volume of 100 µL per sample.

Flash-frozen cells were thawed on ice and resuspended in 1.2 mL IP lysis buffer supplemented with 1× Complete Protease Inhibitor Cocktail (Roche) and 0.1 U µL^−1^ SUPERase•In RNase Inhibitor (Thermo Fisher Scientific). Cells were lysed by sonication and lysates were cleared by centrifugation at 21,000*g* for 15 min at 4 °C. Supernatants were transferred to fresh tubes and protein concentrations were quantified by Bradford assay. For each sample, 0.9 mg of protein was combined with 100 µL of antibody-conjugated beads (prepared as described above) and brought to a final volume of 1.2 mL with supplemented IP–MS lysis buffer. Samples were rotated overnight at 4 °C. The next day, each sample was washed twice with 1 mL of IP–MS wash buffer 1 (50 mM Tris-HCl pH 7.5 at 25 °C, 150 mM NaCl, 5% glycerol, 0.02% Triton X-100) and twice with 1 mL of IP–MS wash buffer 2 (50 mM Tris-HCl pH 7.5 at 25 °C, 150 mM NaCl, 5% glycerol). Beads were then resuspended in 80 µL of elution buffer (50 mM Tris-HCl pH 7.5 at 25 °C, 2 M urea, 1 mM DTT with 0.5 µg mL^−1^ sequencing grade modified trypsin (Promega)) and agitated at room temperature for 1 hour. The supernatants were taken into a new tube. Then beads were resuspended two more times in 60 µL Tris-urea buffer without trypsin and supernatants were combined with the elution. Finally, the supernatants were spun at 5,000*g* for 1 min to pellet any residual beads, and the resulting supernatant was flash-frozen in liquid nitrogen and stored at −80 °C prior to further processing.

For MS sample preparation, first, immunopurified eluates were reduced with 5 mM DTT for 45 min at 25 °C with shaking (600 rpm), then alkylated with 10 mM iodoacetamide (IAA) in the dark for 45 min at 25 °C with shaking (600 rpm). Samples were digested overnight with 0.5 µg sequencing-grade modified trypsin (Promega, V5111) at 25 °C and 600 rpm. Digested peptides were acidified with formic acid, desalted on in-house C18 StageTips (two plugs), dried in a Thermo Savant SpeedVac, and reconstituted in 3% acetonitrile/0.2% formic acid.

For LC-MS/MS analysis, digested peptides were analyzed on a Waters M-Class UPLC fitted with a 15 cm × 75 µm IonOpticks C18 1.7 µm column, coupled to a Thermo Fisher Scientific Q Exactive HF Orbitrap mass spectrometer. Peptides were separated at 400 nl min^−1^ over a 90-min gradient (including sample loading and column equilibration) using Solvent A (0.1% formic acid in water) and Solvent B (0.1% formic acid in acetonitrile). The gradient was: 2% B for 1 min; linear ramp to 10% B over 29 min; to 22% B over 27 min; to 30% B over 5 min; to 60% B over 4 min; to 90% B over 1 min; hold for 2 min; ramp down to 50% B over 1 min, hold for 5 min; ramp down to 2% B over 1 min; and re-equilibration at 2% B for 14 min. Data were collected in data-dependent mode with Xcalibur (4.5.474.0), selecting the 12 most intense peaks per cycle for MS2. MS1 spectra were acquired at 120,000 resolution with an AGC target of 3 × 10^6^ and a scan range of 300–1,800 m/z. MS2 spectra were acquired at 15,000 resolution with an AGC target of 1 × 10^5^, a scan range of 200–2,000 m/z, and an isolation window of 1.6 m/z. Raw files were searched with MaxQuant^59^ (v2.0.3.0) against a combined reference proteome containing the *E. coli* K-12 MG1655 proteome (NCBI RefSeq assembly GCF_000005845.2), the λ-vir proteome (NCBI RefSeq assembly GCF_000840245.1), and the plasmid-encoded DRT3 RT sequences (with an internal 3×FLAG tag on DRT3b). MaxQuant was run with default settings, with the label minimum ratio count set to 1; label-free quantification, iBAQ, and “match between runs” were enabled. One or more unique/razor peptides were required for protein identification. All of the subsequent analyses were performed in python.

For protein comparisons to identify DRT3 protein-interactors, protein groups flagged by MaxQuant as reverse hits, potential contaminants, or identified only by modification site were removed. Zero LFQ intensity values were replaced with 1 before log_2_ transformation. log_2_[fold change] and p-values (unpaired two-tailed t-test) were calculated between WT and untagged control immunoprecipitates, with Benjamini–Hochberg correction applied to control the false discovery rate. Significantly enriched interactors were defined by log_2_[fold change] and adjusted p-value cutoffs as indicated in the figure legend, and visualized as volcano plots in matplotlib. Three independent biological replicates were included per condition.

##### *Eco*DRT3 RT and ncRNA purification

For the purification of *Eco*DRT3 RTs, the genes encoding *Eco*DRT3a (residues 1–398) and *Eco*DRT3b (residues 1–667) were amplified by PCR and cloned into a modified pE-SUMO vector (Life-Sensors) with an N-terminal His_6_-SUMO tag. Mutations were introduced using a PCR-based method, and the sequences were verified by DNA sequencing. For protein expression, *E. coli* Rosetta2 (DE3) (Novagen) cells transformed with either a plasmid encoding His_6_-SUMO-DRT3a or His_6_-SUMO-DRT3b were cultured at 37 °C until the OD_600_ reached 0.8, and protein expression was induced with 0.2 mM IPTG (Nacalai Tesque). The *E. coli* cells were further cultured at 18 °C overnight, harvested by centrifugation, resuspended in buffer A (20 mM Tris-HCl, pH 8.0, 1 M NaCl, 20 mM imidazole, 3 mM 2-mercaptoethanol, and 1 mM PMSF), and lysed by sonication. The lysates were centrifuged, and the supernatant was applied to Ni-NTA Superflow resin (QIAGEN). The bound proteins were eluted with buffer B (20 mM Tris-HCl, pH 8.0, 0.3 M NaCl, 300 mM imidazole, and 3 mM 2-mercaptoethanol). The eluted proteins were incubated with SUMO protease at 4 °C for 3 h and then loaded onto a HiTrap Heparin HP column (Cytiva) equilibrated with buffer C (20 mM HEPES-HCl, pH 7.5, 0.3 M NaCl, and 1 mM DTT). The proteins were eluted using a linear gradient of 0.3–2 M NaCl. For DRT3a, the protein was further purified on a Superdex 200 Increase 10/300 GL column (Cytiva) equilibrated with buffer D (20 mM HEPES-NaOH, pH 7.5, 1 M NaCl, and 1 mM DTT). The purified proteins were stored at −80 °C until use.

The *Eco*DRT3 ncRNA was transcribed *in vitro* with T7 RNA polymerase and purified by 8% denaturing (7 M urea) PAGE. The purity of the RNAs was assessed by denaturing urea–PAGE, with the gel stained with SYBR Gold (Thermo Fisher Scientific).

##### Biochemical DNA synthesis assays

Reverse transcription reactions generally contained *Eco*DRT3a, *Eco*DRT3b, ncRNA, and nucleotide substrates in polymerization buffer (20 mM HEPES, pH 7.5 at 25 °C, 0.3 M NaCl, 5 mM MgCl_2_, and 1 mM DTT). Unless otherwise indicated in the figure legend, 1 mM of total dNTPs, 1 µM of total protein, and 200 nM ncRNA were used in all *in vitro* reverse transcription reactions. Reactions were incubated at 37 °C for 60 min, unless otherwise stated. Radioactive experiments contained trace [α–^32^P]dGTP for *Eco*DRT3a or [α–^32^P]dATP for *Eco*DRT3b, together with unlabelled (cold) dNTPs. The reactions were quenched at 95 °C for 2 min and treated with various nuclease or proteinase reagents before electrophoretic separation, as indicated in figure legends.

Reactions investigating the fate of DNA products were analysed by denaturing urea–PAGE. Unlabelled DNA products were analysed using 15% denaturing urea–PAGE, whereas radiolabelled DNA products were analysed using 5–12% denaturing urea–PAGE. The samples were mixed in equal volumes with 2× RNA Loading Dye (95% formamide, 0.025% Bromophenol blue, and 0.025% SDS), denatured by heating at 95 °C for 5 min, and then loaded onto gels for electrophoretic separation. Non-radioactive gels were stained using SYBR Gold (Thermo Fisher Scientific). Radioactive gels were exposed to a phosphor screen for 16 h at room temperature and imaged using the Typhoon imaging system (GE HealthCare).

Radiolabeled reactions investigating the fate of the *Eco*DRT3-encoded DRT3b enzyme were analysed using 12% Criterion XT Bis-Tris (Bio-Rad Laboratories). The samples were mixed with 6× SDS loading dye before denaturation by heating and electrophoretic separation.

##### *In vitro* biochemical reverse transcription and sequencing

Biochemical DNA synthesis reactions were performed according to the conditions described above, in 20 µL reactions. After quenching the reaction with 95 °C denaturation for 2 min, 1 µg of Proteinase K (Thermo Fisher Scientific) was added, and reactions were incubated at 55 °C for 30 min before heat denaturation. Then, 1 µg of RNase A (Thermo Fisher Scientific) was added to the samples and reactions were incubated at 37 °C for 30 min before heat denaturation. In parallel, three 150-bp oligonucleotides were chemically synthesized (IDT) and pooled at 100 pmol each, then 5′-phosphorylated with T4 PNK (NEB) for 30 min at 37 °C. The oligonucleotide reaction was purified using the Monarch Spin PCR and DNA Cleanup Kit (NEB) and eluted in 20 µL of water. 10 µL of each vitro biochemical reaction was then spiked with 10 pmol of a mixture of three 5′-phosphorylated oligonucleotides (**Table S4**) and cleaned up with the Monarch Spin PCR and DNA Cleanup kit (NEB), according to the Oligonucleotide Cleanup protocol. Libraries were prepared as described above for cDIP and sequenced on the Element AVITI platform in single- or paired-end mode at 75 or 150 cycles per end.

##### Cryo-EM sample preparation and data collection

The purified His -SUMO-DRT3b (0.25 mg mL^−1^) was incubated with 200 μM ddATP and 200 μM ddCTP at 37 °C for 5 min. The mixture (3 μL) was applied to freshly glow-discharged Quantifoil R1.2/1.3 Cu 300-mesh grids covered with a 2 nm carbon film, using a Vitrobot Mark IV (FEI) at 4 °C, with a waiting time of 10 s and a blotting time of 6 s under 100% humidity conditions. The grids were plunge-frozen in liquid ethane cooled at liquid nitrogen temperature. Cryo-EM data were collected using a Titan Krios G3i microscope (Thermo Fisher Scientific), running at 300 kV and equipped with a Gatan Quantum-LS Energy Filter (GIF) and a Gatan K3 Summit direct electron detector in correlated double sampling mode (The University of Tokyo, Japan). Movies were recorded at a nominal magnification of 105,000×, corresponding to a calibrated pixel size of 0.83 Å, with an electron exposure of 7.77 e^−^ pix^−1^ sec^−1^ for 4.66 s, resulting in an accumulated exposure of 50.6 e^−^ Å^−2^. The data were automatically acquired by the image shift method using the EPU software (Thermo Fisher Scientific), with a defocus range of −0.8 to −2.0 μm and 835 movies were acquired.

##### Image processing

Data processing was performed using the cryoSPARC v5.0.2 software package^60^. Dose-fractionated movies were aligned using the Patch motion correction, and the contrast transfer function (CTF) parameters were approximated using Patch-Based CTF estimation with the default settings. Particles were automatically picked using Topaz Picker^61^. The picked particles were subjected to 2D classification, and 13,084 selected particles were used for Ab initio Reconstruction imposed. The resulting map was used as an initial template for Homogeneous Refinement with D3 symmetry and the default parameters. Following Reference-Based Motion Correction, the particles were further refined using Non-uniform Refinement^62^. This refinement yielded a map at 3.1-Å resolution according to the gold-standard Fourier shell correlation (FSC) = 0.143 criterion^63^. Local resolution was estimated using BlocRes in cryoSPARC^64^.

##### Model building and validation

The models of the DRT3b hexamer were built using COOT^65^, starting from a model predicted by Boltz-2^66^. The models were refined using Servalcat^67^ against unsharpened half maps, with structural restraints derived from Boltz-2 predictions processed by ProSMART^68^. The models were validated using MolProbity^69^. The statistics of the 3D reconstruction and model refinement are summarized in **Table S5**. Molecular graphics and cryo-EM density map figures were prepared using CueMol (http://www.cuemol.org) or UCSF ChimeraX^70^.

##### Conservation of DRT3b priming and templating residues

The seven C-terminal residues of the RT sequences, along with the five residues spanning E22 and R241 of DRT3b, were extracted from the UG/RT tree of a previous study^3^ and aligned using the DECIPHER^71^ (v3.8.0) package in R. Alignment positions with occupancy below 50% were trimmed, and the resulting alignments were visualized with ggplot2 and used to generate a WebLogo^72^ (v3.7.9).

##### Escaper phage isolation

To isolate phages that resist DRT3-mediated defense, serial dilutions of λ-vir were mixed in soft molten agar containing *E. coli* K-12 strain MG1655 with plasmids encoding for DRT3 or an EV control and plated onto LB agar plates with chloramphenicol (25 µg mL^−1^). After overnight incubation at 37 °C, the plates were removed and plaques were picked into 1 mL of the same host strain culture (grown to OD_600_ of 0.1) used for plating. Cultures were incubated with shaking at 37 °C for 3 h, after which 800 µL of each was transferred to a fresh tube and treated with chloroform (5% v/v final) to lyse residual cells. After centrifugation at 4,000*g* for 5 min to separate the aqueous phase from debris and chloroform, 5 µL of the spun down supernatant was taken into added to a fresh culture of *E. coli* cells (OD_600_ of 0.1) and propagated shaking with shaking at 37 °C for 3 h. Amplification was repeated for three rounds total before confirming escaper phages by small-drop plaque assay.

##### Escaper phage whole-genome sequencing

Genomic DNA from escaper phages were isolated by treating 88 µL of phage in LB with 0.2 U of DNase I (NEB) and 10 µg of RNase A (Thermo Fisher Scientific) in 1 × DNase buffer (10 mM Tris-HCl pH 7.5 at 25 °C, 2.5 mM MgCl_2_, 0.5 mM CaCl_2_). After incubation at 37 °C for 30 min, the enzymes were heat inactivated at 75 °C for 10 min. Next, the phage was lysed in phage lysis buffer (10 mM Tris-HCl, 10 mM EDTA, 0.5% SDS) and 20 µg of Proteinase K (Thermo Fisher Scientific) at 55 °C for 30 min. Following lysis, DNA was purified with the Genomic DNA Clean & Concentrator-25 kit (Zymo).

Genomic DNA was sequenced using a previously described method^73^. Briefly, phage genomic DNA was tagmented using purified in-house TnY. 10 ng of purified genomic DNA (gDNA) was tagmented with TnY previously incubated with Nextera read 1 and read 2 oligos (**Table S4**). Following incubation at 55 °C for 7 minutes, the reaction was treated with 16 U mL^−1^ Proteinase K (NEB) and cleaned up with the Monarch Spin PCR and DNA Cleanup kit (NEB) kit. PCR amplification and Illumina barcoding was done for 20 cycles with KAPA HiFi Hotstart ReadyMix, with an annealing temperature of 63 °C and an extension time of 1 min. The PCR reactions were pooled and resolved on a gel. The 400–800 bp smear was gel extracted with the Qiaquick Gel Extraction kit (Qiagen) and sequenced on the Element AVITI in paired-end mode with 150 cycles per end.

The resulting datasets were processed using Cutadapt^50^ (v.5.0) to remove adapter sequences, trim low-quality ends from reads and filter out reads shorter than 15 bp. Reads were mapped to the λ-vir (NCBI: NC_001416.1) genome using Bowtie2^73^ (v.2.2.1) with the default parameters. SAMtools^52^ (v.1.17) was used to sort and index alignments, and coverage tracks were visualized in IGV^54^ (v.2.17.4). Mutations were identified with breseq^75^ (v.0.39.0) using alignments from Bowtie2 and the same λ-vir reference genome as above. λ-vir escaper mutations are listed in **Table S6**.

##### Infection and trigger response growth curves

Overnight cultures of *E. coli* MG1655 transformed with DRT3^WT^ or DRT3a/b^YVAA^ plasmids were diluted 1:100 in LB supplemented with the appropriate antibiotics. After reaching OD_600_ of 0.2, the cultures were spun down at 4,000*g* for 5 min at room temperature and washed before resuspending in LB with antibiotics. 180 µL of each resuspended cell culture were transferred to individual wells on a 96-well optical plate. For phage infection, 20 µL of each corresponding condition was added to the plate: LB as a mock infection control or λ-vir at a final MOI of 0.5 or 5. For trigger expression, 20 µL of either 10% glucose or 2% arabinose was added for a final concentration of 1% glucose or 0.2% arabinose. The plate was incubated for the times indicated in the figures at 37 °C with shaking. OD_600_ values were recorded every 10 min using the Synergy Neo2 microplate reader (Biotek).

##### Colony-formation time course analysis

*E. coli* MG1655 was transformed with one of two pBAD plasmids (pBAD-EV or pBAD-Gam) and one of two DRT3 plasmids (DRT3WT or DRT3a/b^YVAA^). Overnight cultures of each combination were diluted 1:100 in LB supplemented with 1% glucose and the appropriate antibiotics. After reaching an OD_600_ of 0.2, the cultures were spun down at 4,000*g* for 5 min at room temperature, washed with 1× TBS, and resuspended in LB with antibiotics. Each culture was split, and arabinose was added to a final concentration of 0.2% for 30, 60, or 90 min before quenching with 1% glucose. Cells were then serially diluted in LB containing 1% glucose and spotted onto LB agar plates with 1% glucose and antibiotics, and growth phenotypes were assessed after overnight incubation at 37 °C.

##### Cell viability assays with trigger and host factor candidates

To assess the effects of DRT3 and its catalytic mutants co-expression with λ Gam, WT trigger and escaper e5 genes were cloned onto an expression vector under the control of an arabinose-inducible araBAD promoter (pBAD). *E. coli* K-12 strain MG1655 cells expressing either DRT3^WT^ or DRT3a^YVAA^/ DRT3b^WT^ were transformed with trigger plasmids and grown under repressive conditions in LB medium supplemented with 1% glucose. Overnight cultures were centrifuged at 4,000*g* for 5 min and cell pellets were washed with fresh LB to remove residual glucose. Cells were pelleted again as before and resuspended in fresh LB. 20 µL of concentrated transformants were drip plated on LB agar plates supplemented with chloramphenicol (25 µg mL^−1^), spectinomycin (100 µg mL^−1^) and arabinose (0.2%) or glucose (1%). Plates were incubated overnight at 37 °C and imaged the next day.

To assess the cell viability of the various gene deletion mutants described in **Table S2**, knock-out strains were transformed with either DRT3^WT^ or DRT3a/b^YVAA^. To assess synthetic lethality, cells were recovered in 10 mL of LB medium at 37 °C for 4 hours. After recovery, cultures were diluted 1:10 in fresh LB, and 10× serial dilutions were plated on LB agar plates supplemented with chloramphenicol and kanamycin. To assess for phage defense, individual colonies were transformed and used in small-drop plaque assays.

##### Quantification of dsDNA by PicoGreen assay

dsDNA from *in vitro* biochemical transcription reactions were quantified using a PicoGreen-like dye (Apexbio Technology, K1603). Following initiation of the *in vitro* reaction as described above, 5 µL of sample was removed from the total reaction volume at intervals every 0, 5, 10, 15, 20, 40, and 60 mins. These aliquots were immediately quenched by 45 µL of TE (10 mM Tris-HCl, 1 mM EDTA, pH 7.5). After all samples were quenched, each reaction was prepared for quantification via manufacturer instructions in a final volume of 100 µL. The fluorescence signal was read with the Synergy Neo2 microplate reader (Biotek) at Ex = 480 nm, Em= 520 nm.

##### Tn–seq screening for host factors

A vector containing a kanamycin resistance marker within a Himar1 *Mariner* mini-transposon designed for Tn–seq (Addgene 102939) was transformed into an MFDpir *E. coli*, which is a diaminopimelic acid (DAP) auxotrophic conjugative donor. Tn–seq vector containing conjugative donors was selected for, propagated, and banked in DAP (0.3 mM), carbenicillin (100 µg mL^−1^), and kanamycin (50 µg mL^−1^). Tn–seq conjugative donors were then conjugated with MG1655 *E. coli* to generate a *Mariner* Tn–seq library with a MG1655 background. Briefly, donor and recipient cells were grown overnight from frozen glycerol stocks. Donor cultures were diluted 1:20 in 50 mL of fresh LB with DAP, carbenicillin, and kanamycin, while recipient cells were diluted 1:50 in 50 mL of fresh LB. Cultures were grown to OD_600_ 0.4 and washed a total of two times by centrifuging cells at 4,000*g* for 5 minutes at room temperature, resuspending cells in 40 mL of PBS, and repeating once more. After the final wash, each pellet was resuspended in 1 mL of LB. The donor and recipient inocula were mixed in a 1:2 (donor:recipient) ratio, with a cellular count equivalent of 1 mL of OD_600_ 1 for the donor. The donor:recipient conjugation mixtures were placed onto sterile cellulose filters on DAP-containing LB plates. The conjugation mixture was incubated for 3 hours at 37 °C. The filters were then removed, and the mixture was resuspended in 45 mL of LB using a vortex. The resuspended cells were then centrifuged at 4,000*g* for 5 minutes at room temperature. The pellets were resuspended in 1 mL of LB and then plated on large kanamycin-containing LB bioassay plates. The plates were grown overnight at 37 °C for 18 hours, or until there was enough biomass and individual colonies were still identifiable. The cells were then scraped off into 10 mL of LB, which was then split into 500 μL heavy inoculum glycerol shots. This was repeated a total of five times for replicate library generation. To screen for host factors, each independent replicate Tn–seq MG1655 *E. coli* library was made electrocompetent. One of two single effector plasmids containing both DRT3 and pBAD-Gam (DRT3^WT^ or DRT3a/b^YVAA^) was transformed into each library and recovered for 2 hours in SOC recovery media supplemented with 1% glucose. The transformation mixture was diluted 1:10, and then 500 μL was plated onto large LB bioassay plates containing either 1% glucose or 0.2% arabinose and the appropriate antibiotics. These plates were then grown at 37 °C for 18 hours, after which all the colonies were scraped off. gDNA from these cells was extracted using the Wizard Genomic DNA Purification kit (Promega). Extracted gDNA was digested with MmeI (2U MmeI/1 μg of gDNA) at 37 °C for 1 hour, followed by inactivation at 65 °C for 20 minutes. This was then purified using QIAquick PCR purification columns. Sequencing adaptors were then ligated onto digested mini-transposons with T4 ligase at 16 °C for 14 hours. After another QIAquick PCR purification cleanup, PCR amplification and Illumina barcoding were done for a total of 25 cycles with Q5 Hot Start HF DNA polymerase, with an annealing temperature of 65 °C and an extension time of 20 seconds. The 173 bp band was gel extracted with the Qiaquick Gel Extraction kit (Qiagen) and sequenced on the Element AVITI in single-end mode.

Raw reads were processed with bbduk^76^ (v39.83) to filter for Himar transposon-end containing reads, to trim adapter sequences, low-quality bases, and to trim transposon sequence to leave only the 16-17 bp of genomic sequence that can be mapped back to the *E. coli* MG1655 reference genome (NC_000913.3). Processed reads were then aligned with Bowtie2^73^ (v1.3.1) with no allowance for mismatches. Alignments were sorted and indexed using SAMtools^52^ (v1.17). Strand-specific coverage tracks were generated with bamCoverage^53^ (v3.5.6) with no normalization metrics and parameters that identified the TA within the read as the insertion site (bin size == 1 and offset == −1). Coverage tracks were visualized in IGV^54^. Mapped reads were assigned to annotated genes with featureCounts^55^ (v.2.0.2) (--read2pos 5). The resulting count matrices were passed to PyDESeq2^56^ (v.0.5.3) for differential insertion analysis using default parameters identical to the RNA–seq analysis above. Results were displayed as volcano plots generated in matplotlib, with log_2_[fold change] plotted as a function of −log_10_[p_adj_]. Five independent biological replicates were included for every comparison.

##### Functional enrichment analysis (GSEA)

To categorize host genes differentially regulated by active DRT3 during λ-vir infection, Gene Set Enrichment Analysis (GSEA) was performed on the host transcripts of RNA–seq data comparing DRT^WT^ + λ infection to DRT3a/DRT3b^YVAA^ + λ infection. Differential expression was first computed using Py-DESeq2^56^ (v.0.5.3) on host-only count matrices, and genes were ranked in descending order by log_2_ [fold change].

GO Biological Process gene sets for *E. coli* K-12 MG1655 (NCBI taxonomy ID 511145) were constructed from the NCBI gene_info and gene2go annotation files, and only annotations in the Process subontology were kept. Gene IDs were mapped to gene symbols, and gene sets containing fewer than 5 genes were discarded, leading to 454 total gene sets for analysis. Preranked GSEA was performed using gseapy^77^ (v1.1.13) with the following parameters: minimum gene set size 5, maximum gene set size 500, 1,000 permutations, and a fixed random seed (42) for reproducibility. Significantly enriched pathways had a Benjamini–Hochberg FDR q-value below 0.05 (p_adj_). Results were graphically visualized as a bubble plot with NES on the x-axis and bubble size scaled to −log_10_(p_adj_). Significantly enriched (NES > 0) and depleted (NES < 0) pathways were plotted with matplotlib (v3.10.5).

##### Nucleotide quantification by LC–MS/MS

Metabolomics data were collected and analyzed as previously described^9^. *E. coli* K-12 strain MG1655 cells were transformed with DRT3^WT^ or DRT3a/b^YVAA^ plasmids. Individual colonies were inoculated into LB medium supplemented with chloramphenicol (25 µg mL^−1^) and grown overnight at 37 °C with shaking. The next day, cultures were diluted 1:100 in 20 mL fresh LB medium and grown to OD_600_ of 0.3. Cultures were split in half and λ-vir phage was added to one half at an MOI of 5. Uninfected and infected cultures were grown for an additional 30 min at 37 °C with shaking. Cells were centrifuged at 4,000*g* for 5 min at 4 °C and pellets were washed with 1 mL of cold 1× TBS. Cells were centrifuged again at 15,000*g* for 1 min at 4 °C. After supernatants were removed, pellets were flash-frozen in liquid nitrogen and stored at −80 °C.

Cell lysates were prepared as previously described^78^. Flash-frozen pellets were thawed on ice and resuspended in 600 µL of 100 mM sodium phosphate buffer (10:1 mixture of Na_2_HPO_4_ and NaH_2_PO_4_) supplemented with 1 mg mL^−1^ lysozyme. Cells were mechanically lysed using a bead beater (2.5 min × 2 cycles at 4 °C). Lysates were centrifuged at 4,000*g* for 15 min at 4 °C and supernatants were transferred to new tubes. The samples were transferred to Amicon Ultra-0.5 3 kDa filter units (Merck Millipore) and centrifuged at 12,000*g* for 45 min at 4 °C. Filtrates containing metabolites were collected and stored at −80 °C. After retrieval from −80 °C, and before use for injection in LC–MS/MS, filtrates were additionally purified by centrifugation at 20,000*g* for 20 min at 4 °C. To confirm that each condition contained equal amounts of cellular material, the protein concentration in filtration supernatants was measured using the BCA assay (Thermo Fisher Scientific). Protein concentrations varied by less than 10% across all samples.

Nucleotide and deoxynucleotide profiling analysis was carried out with ion-paired reversed-phase liquid chromatography using an HPLC (1290 Infinity II, Agilent Technologies) system coupled to a triple quadrupole mass spectrometer (6495D, Agilent Technologies) with electrospray ionization operated in negative mode. The column was a ZORBAX RRHD Extend-C18 (2.1 × 150 mm, 1.8 µm pore size; 759700-902, Agilent Technologies). Then, 5 µL of the experimental samples was injected. Mobile phase A was 3% methanol (in H_2_O), and mobile phase B was 100% methanol. Both mobile phases contained 10 mM of the ion-pairing agent tributylamine (90780, Sigma-Aldrich), 15 mM acetic acid, and 5 µM medronic acid (5191-4506, Agilent Technologies). The LC gradient was: 0–2.5 min 100% A, 2.5–7.5 min ramp to 80% A, 7.5–13 min ramp to 55% A, 13–20 min ramp to 99% B, 20–24 min hold at 99% B. The flow rate was 0.25 mL min^−1^, and the column compartment was heated to 35 °C. The column was then backflushed with 100% acetonitrile (0.25 mL min^−1^ flow rate 24.05–27 min, followed by 0.8 mL min^−1^ flow rate 27.5–31.35 min and 0.6 mL min^−1^ flow rate, 31.35–31.50 min) and re-equilibrated with 100% A (0.4 mL min^−1^ flow rate 32.25–40 min). The conditions of the MRM transitions for dADP, dATP, dTDP and dTTP were as follows (collision energy (V)): dADP, 410 > 158.7 (24), 410 > 79.1 (44); dATP, 490 > 391.9 (24), 490 > 158.9 (32); dTDP, 401 > 97 (25), 401 > 79 (44); dTTP, 481 > 383.1 (20), 481 > 158.8 (36). Fragmentor and CAV were kept constant at 166 V and 4 V, respectively. For other nucleotides and deoxynucleotides measured, conditions of the MRM transitions are shown in **Table S7**. Moreover, the identity of each compound was confirmed with subsequent injection and acquisition of a pure chemical standard. Data analysis, including peak area integration and signal extraction, was performed with Skyline-daily^79^ (v.24.1.1.398).

##### Quantification and statistical analysis

Phage defense was quantified as the fold-reduction in efficiency of plating (EOP) relative to the empty-vector control, from 3 independent technical replicates unless otherwise indicated in the figure legend. Growth curves, colony-formation assays, and PicoGreen time courses were performed in 3 independent biological replicates. Data are shown as mean ± SD.

Covariation in the ncRNA covariance model was evaluated with R-scape^49^ (v2.6.10) at an E-value threshold of 0.05.

Differential expression analysis of total RNA–seq data and differential transposon-insertion analysis of Tn–seq data were performed with PyDESeq2^56^ (v0.5.3) using the Wald test with Benjamini-Hochberg correction to control the false discovery rate. RNA–seq comparisons included three biological replicates and Tn-seq comparisons included five biological replicates. Results were visualized as volcano plots of log_2_ [fold change] versus –log_10_ [p_adj_]. Gene Set Enrichment Analysis was performed with gseapy^77^ (v1.1.13) in preranked mode on genes ranked by log_2_[fold change], using 1,000 permutations and a fixed random seed (42). GO Biological Process-derived pathways with a Benjamini-Hochberg q-value < 0.05 were considered significantly enriched and reported as normalized enrichment scores (NES).

For co-immunoprecipitation mass spectrometry, log_2_[fold change] and p-values were calculated between DRT3^WT^ and DRT3a/b^YVAA^ conditions using an unpaired two-tailed t-test with Benjamini-Hochberg correction. Enriched protein interactions were defined by the log_2_[fold change] > 3.32 and adjusted-p cutoffs < 0.05, across three independent biological replicates.

Nucleotide and deoxynucleotide pools were quantified by LC–MS/MS (Skyline-daily^78^ v24.1.1.398), with sample loading normalized by BCA assay (<10% protein variation across conditions) from three independent biological replicates.

Cryo-EM resolution was based upon FSC = 0.143 criterion. Refinement and validation statistics are summarized in **Table S5**.

## SUPPLEMENTAL VIDEO AND EXCEL TABLE TITLES AND LEGENDS

**Video S1 | Structure of the DRT3b hexameric complex, Related to Figure 3**.

The DRT3b protomers are shown as ribbon and surface models, while the DNA product is depicted as a cartoon model.

**Video S2 | Structure of the DRT3b monomeric complex, Related to Figure 3**.

The DRT3b protomer is shown in ribbon and surface models, while the DNA product is depicted as a cartoon model, with disordered regions indicated by dotted lines. E22, R241, and Y666 are represented as stick models, and a Mg^2+^ ion is depicted as an orange sphere.

**Video S3 | Structure of the DRT3b active site, Related to Figure 3**.

DNA-contacting residues, including E22 and R241, are highlighted as thick stick models, with cryo-EM density shown as a transparent gray surface.

**Table S1 | Description and sequence of plasmids used in this study, Related to STAR Methods.**

**Table S3 | DRT3 proteins and RNAs tested in this study, Related to STAR Methods.**

**Table S7 | Conditions of MRM transitions from metabolomics measurements, Related to Figure 5**.

