## Supplemental Figures and Tables S2, S4, S5, S6 for "Coordinated RNA- and protein-templated synthesis of double-stranded DNA by a dual reverse transcriptase immune system"

### SUPPLEMENTARY INFORMATION

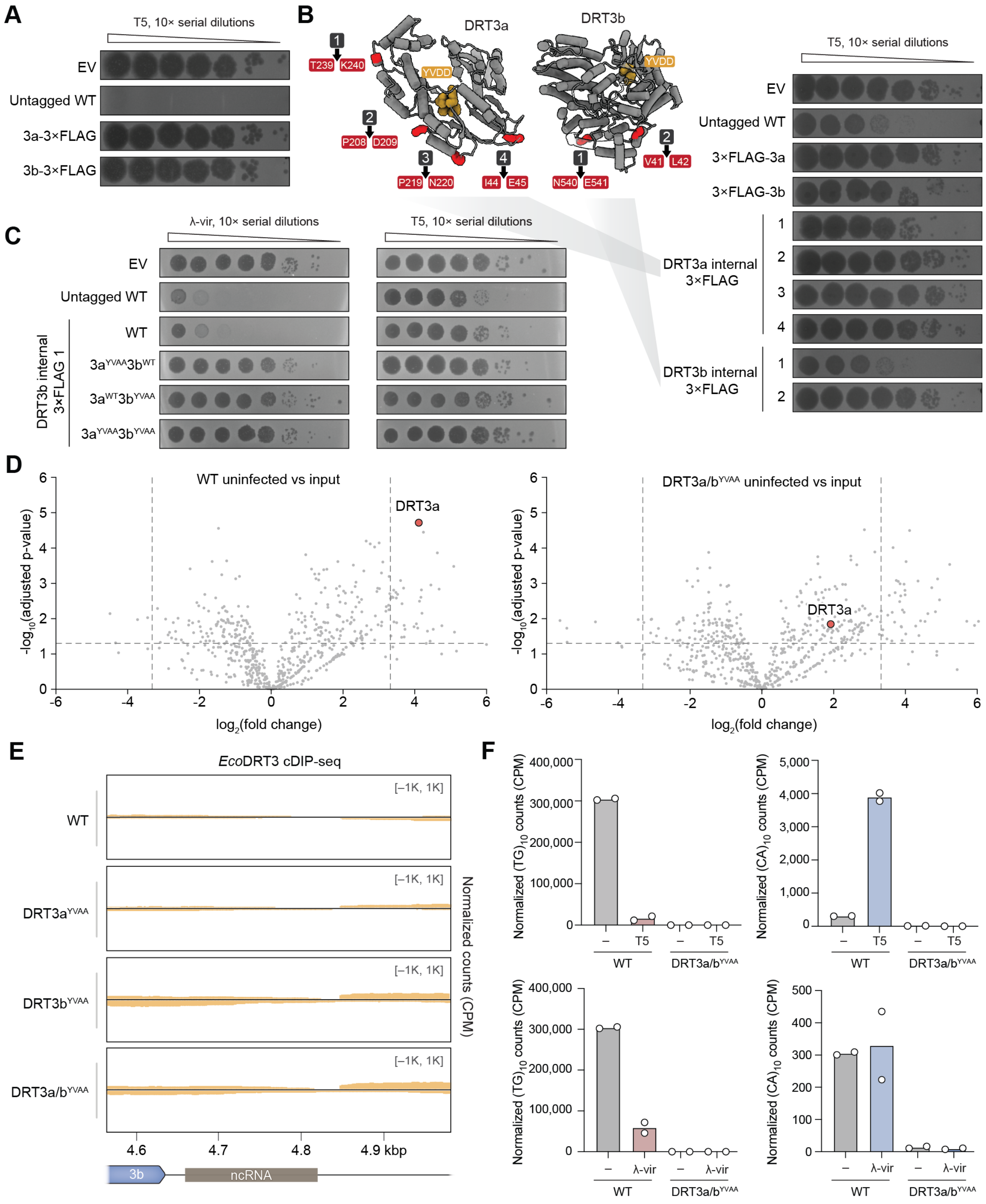

**Figure S1 | Tagging constraints, mutational analysis, and biochemical characterization of *Eco*DRT3, Related to Figure 1.**

**(A)** Plaque assays showing T5 phage resistance for empty vector (EV), untagged WT *Eco*DRT3, and C-terminal 3×FLAG tags. Terminal tagging abolishes defense activity.

**(B)** AlphaFold3 (AF3) structure predictions of *Eco*DRT3a and *Eco*DRT3b highlighting candidate loop regions selected for internal 3×FLAG insertion. Right, plaque assays of N-terminal 3×FLAG tags and internal 3×FLAG variants in DRT3a and DRT3b showing that a single internal 3×FLAG-tagged DRT3b site retains WT phage defense activity.

**(C)** Plaque assays with λ-vir (left) and T5 (right) phage testing internal 3×FLAG-tagged DRT3b variants and combinations of active site mutants in DRT3a and DRT3b, demonstrating that intact active sites in both enzymes are required for defense.

**(D)** Volcano plots from co-immunoprecipitation followed by mass spectrometry (co-IP MS) experiments using 3×FLAG-tagged DRT3b, showing enrichment of DRT3a in WT uninfected cells (left) that is reduced in catalytic mutant backgrounds (right), indicating that DRT3a–DRT3b complex formation depends on intact RT active sites. Dashed lines indicate thresholds for statistical significance ($p_{adj} < 0.05$) and fold-change cutoffs ($|\log_2 FC| > 3.32$).
**(E)** Genome browser tracks of cDIP-seq signal across the DRT3 locus for uninfected WT and RT mutant strains, with the coordinates of the ncRNA displayed below the x-axis.
**(F)** Quantification of normalized cDIP-seq reads containing $(dTdG)_{10}$ (left) and $(dCdA)_{10}$ (right) motifs, plotted as counts per million (CPM), for WT and DRT3a/b catalytic mutant backgrounds under uninfected and T5 phage (top) or λ phage (bottom) infected conditions. Data represent the mean with individual data points from *n* = 2 independent biological replicates.

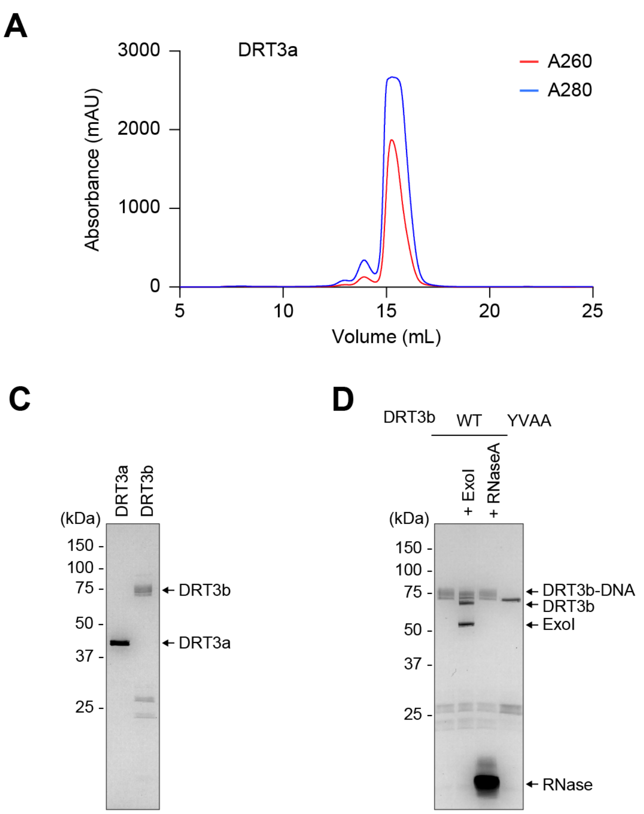


**Figure S2 | Preparation of *Eco*DRT3a and *Eco*DRT3b, Related to Figure 2.**
**(A)** Size-exclusion chromatography profile of *Eco*DRT3a resolved on a Superdex 200 Increase 10/300 GL column.
**(B)** Affinity chromatography profile of *Eco*DRT3b resolved on a HiTrap Heparin HP column. Note the high A260/A280 ratio, indicative of co-purifying nucleic acid.
**(C)** SDS–PAGE of final purified DRT3a and DRT3b proteins used in biochemical and structural studies.
**(D)** SDS–PAGE analysis of purified DRT3b before or after treatment with ExoI and RNase A, as indicated. The diffuse band observed upon purification is partially reduced following ExoI treatment, with the appearance of a faster-migrating band comparable to that observed with the catalytically inactive YVAA mutant. These results are consistent with DRT3b being conjugated to DNA of heterogeneous length during cellular expression and purification.

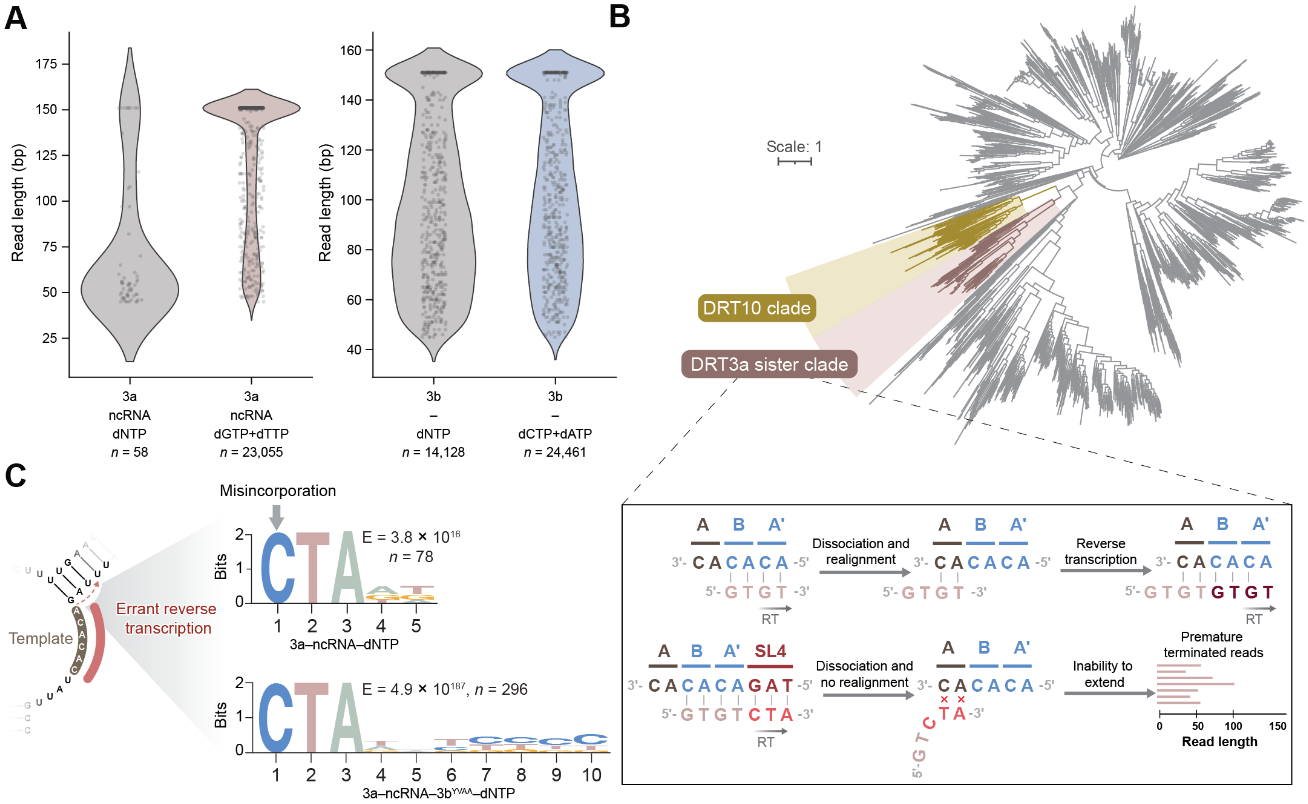


**Figure S3 | Read-length distributions, phylogenetic context, and termination signatures of DRT3 DNA synthesis products, Related to Figure 2.**

**(A)** Violin plots showing length distributions of *in vitro* DNA products generated by DRT3a–ncRNA and DRT3b under the indicated nucleotide conditions. DRT3a produces substantially shorter products in the presence of all four dNTPs compared to reactions containing only dGTP and dTTP, whereas DRT3b product lengths are largely unchanged across conditions. The numbers of reads (n) are indicated below each condition and up to 500 individual data points were plotted per condition.

**(B)** Unrooted phylogenetic tree (top) of representative Unknown Group reverse transcriptases, highlighting the placement of DRT3a within a clade related to DRT10 and other RNA-templated systems. A schematic model (bottom) illustrates the proposed mechanism of template realignment during iterative synthesis, and how readthrough into adjacent stem–loop structures (i.e., SL4) may impair reannealing and processivity.

**(C)** Sequence logos derived from the 3′ termini of unmapped $(dTdG)_{10}$ reads generated in DRT3a–ncRNA (top) and DRT3a–ncRNA–DRT3b$^{YVAA}$ (bottom) reactions containing all four dNTPs, revealing enrichment of non-(dT/dG) nucleotides at terminal positions. This pattern is consistent with misincorporation and extension beyond the canonical template region, leading to premature termination of poly-(dTdG) synthesis. Statistical significance of motif enrichment is indicated (E-value), with total reads analyzed (n) shown.

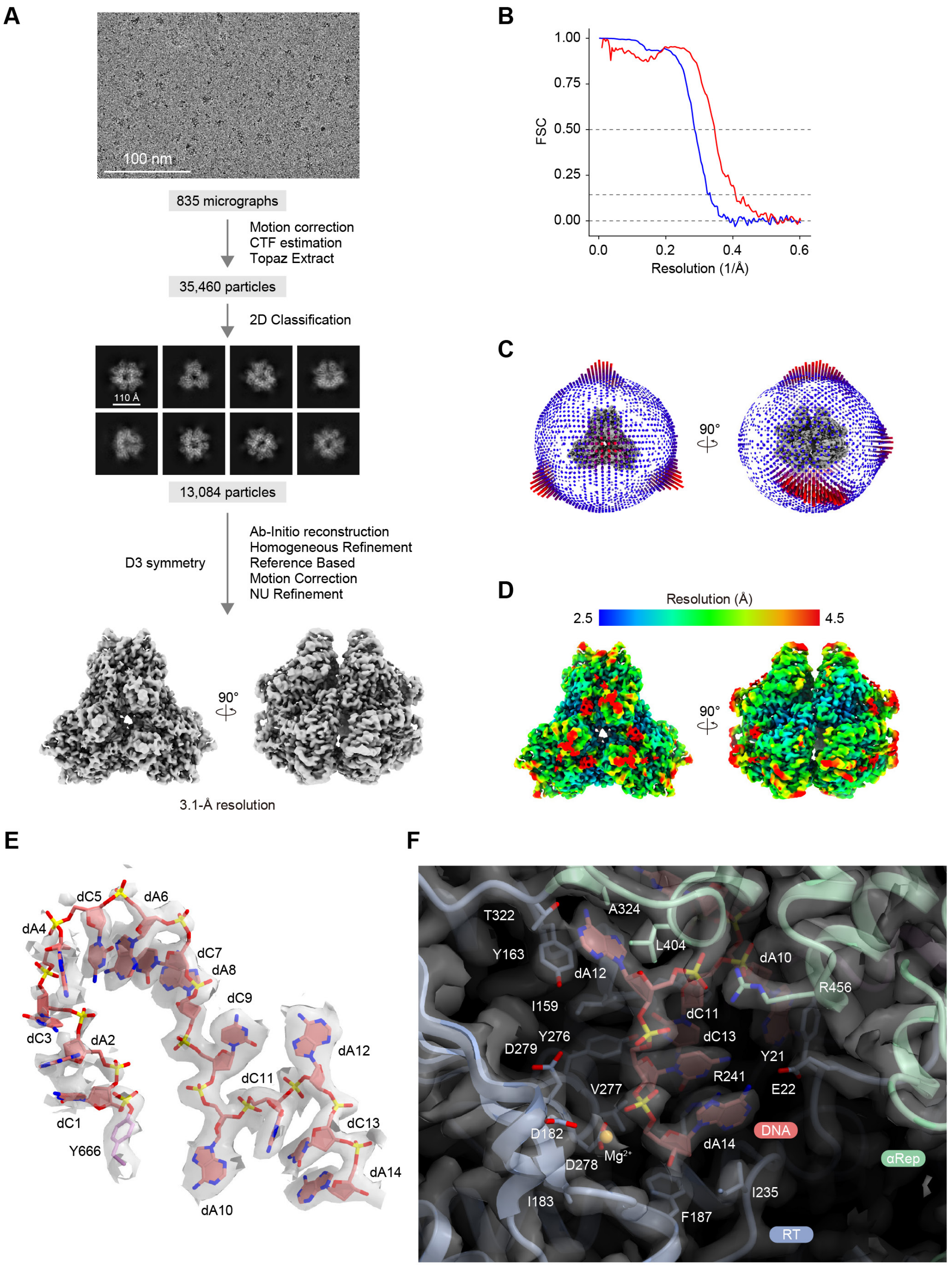


**Figure S4 | Cryo-EM analysis of the DRT3b hexameric complex, Related to Figure 3.**
**(A)** Single-particle cryo-EM image processing workflow.
**(B)** FSC curves. The map-to-map FSC curves were calculated between the two independently refined half-maps after masking (blue line), and the overall resolution was determined by the gold standard FSC = 0.143 criterion. The map-to-model FSC curves were calculated between the refined atomic models and maps (red line).
**(C)** Angular distribution of particles in the final reconstruction.
**(D)** Cryo-EM density maps colored according to the local resolution.
**(E)** Structure of a 14-mer $(dCdA)_7$ strand covalently attached to Y666, along with the cryo-EM density map (semi-transparent gray surface) at a lower contour level than in Figure 3E, illustrating the connection at the flexible dC1–dC5 nucleotides.
**(F)** Cryo-EM density map showing the DRT3b active site.

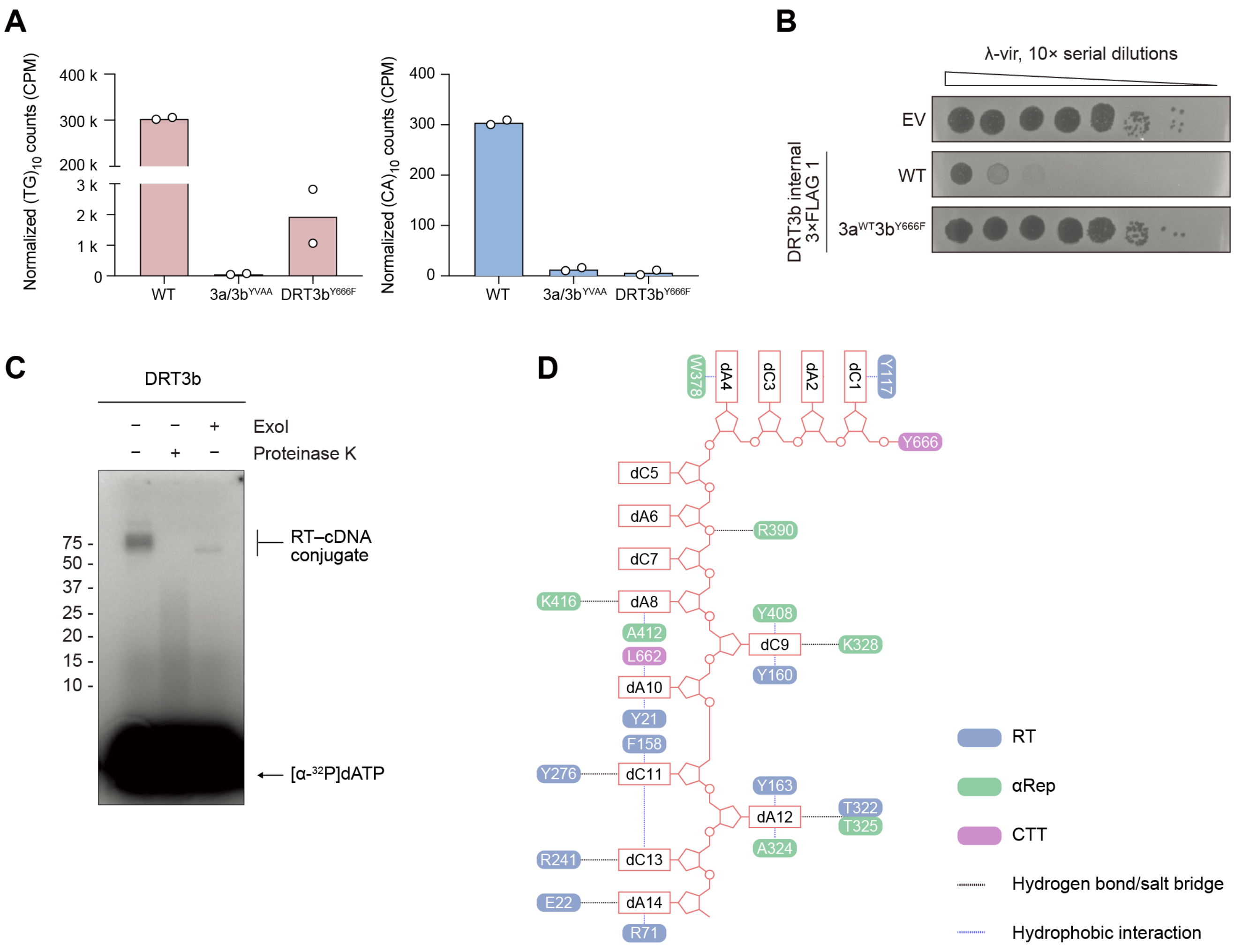


**Figure S5 | Biochemical activity and structural interactions between DRT3b and its DNA product, Related to Figure 3.**
**(A)** Quantification of $(dTdG)_{10}$ and $(dCdA)_{10}$ reads in WT and mutant backgrounds, plotted as counts per million (CPM). Data represent the mean with individual data points from $n = 2$ independent biological replicates.
**(B)** Plaque assays with λ-vir testing the indicated DRT3a$^{WT}$ and DRT3b$^{Y666F}$ mutant, demonstrating the essentiality of priming residue Y666 for phage defense.
**(C)** *In vitro* DNA synthesis by wild-type DRT3b. Reaction products were analyzed by denaturing gel electrophoresis, showing formation of DNA products and disruption of the product band with proteinase K treatment and collapse of the protein–DNA conjugates with exonuclease I treatment.
**(D)** Schematic showing interactions between DRT3b and the DNA product. Residues from distinct domains (RT-like, αRep, and CTT) are color-coded.

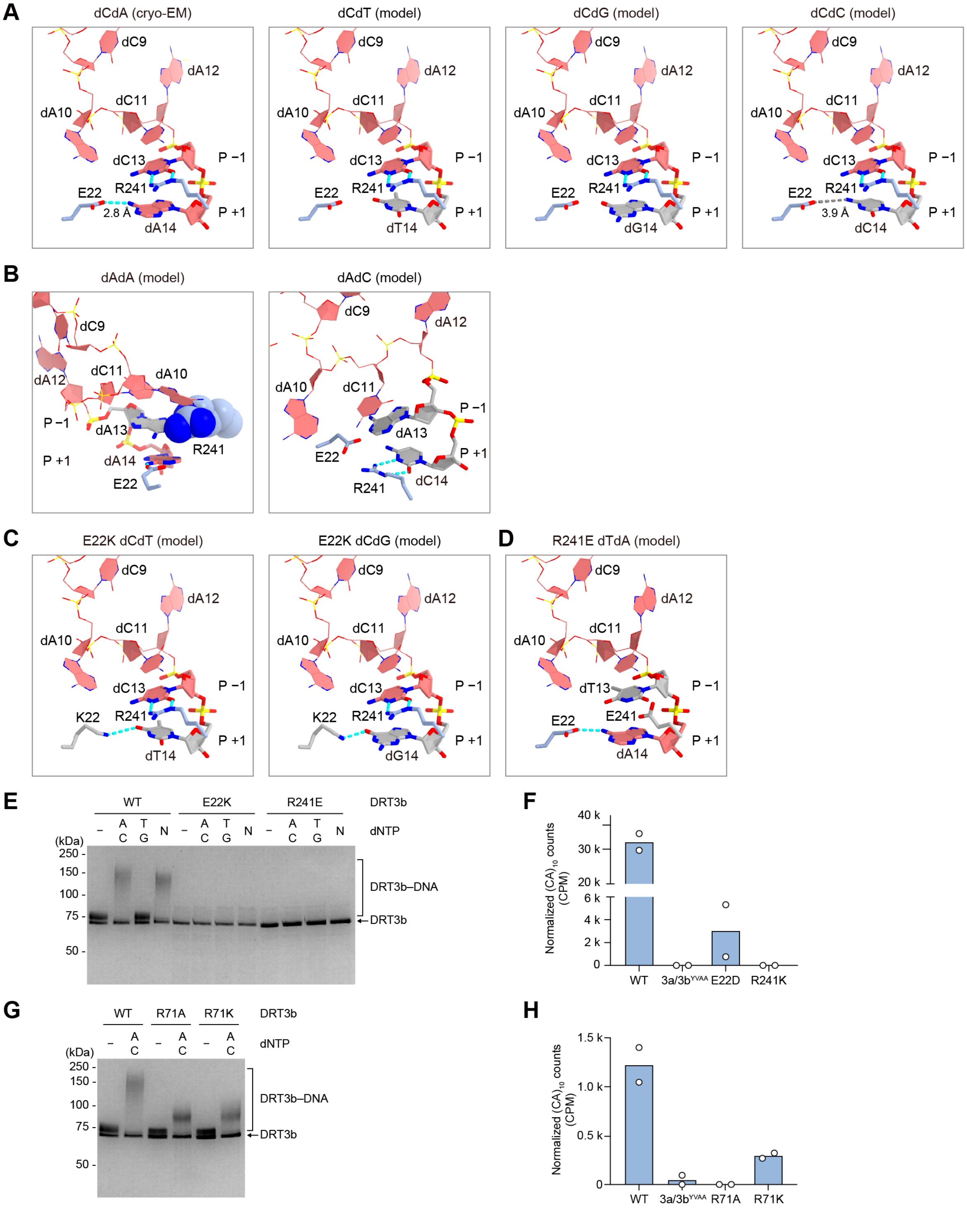


**Figure S6 | Structural basis of nucleotide selection by DRT3b, Related to Figure 3.**
**(A)** Structure of the DRT3b active site and modeling of different nucleotides at position +1. Hydrogen bonds between E22 and dA14 and between R241 and dC13 are depicted with cyan dashed lines. Gray dashed lines denote the absence of a predicted hydrogen bond, as for E22 and dC14 in the dCdC model. The distances between the carboxyl group of E22 and the amino groups of dA14 or dC14 are indicated.
**(B)** Modeling of adenine at position −1 and cytosine at position +1. Adenine at position −1 would sterically clash with the R241 side chain (depicted as a space-filling model), suggesting that when adenine occupies position −1, the R241 side chain would move from position −1 to position +1, where it interacts with dCTP.
**(C)** Modeling of thymine and guanine at position +1 in the E22K mutant.
**(D)** Modeling of thymine at position −1 in the R241E mutant.

**(E)** SDS–PAGE analysis of DRT3b DNA synthesis reactions in the presence of dATP and dCTP, dTTP and dGTP, or all four dNTPs.
**(F)** Quantification of $(dCdA)_{10}$ reads in WT, DRT3a/b$^{YVAA}$, 3b$^{E22D}$, and 3b$^{R241K}$ mutant backgrounds in *E. coli* cells, plotted as counts per million (CPM). Data represent the mean of $n = 2$ independent biological replicates.
**(G)** SDS–PAGE analysis of DRT3b DNA synthesis reactions in the presence of dATP and dCTP, showing a reduction in the covalent DRT3b–DNA adduct in the presence of R71A or R71K mutations.
**(H)** Quantification of $(dCdA)_{10}$ reads in WT, DRT3a/b$^{YVAA}$, 3b$^{R71A}$, and 3b$^{R71K}$ mutant backgrounds in *E. coli* cells, plotted as counts per million (CPM). Data represent the mean of $n = 2$ independent biological replicates.

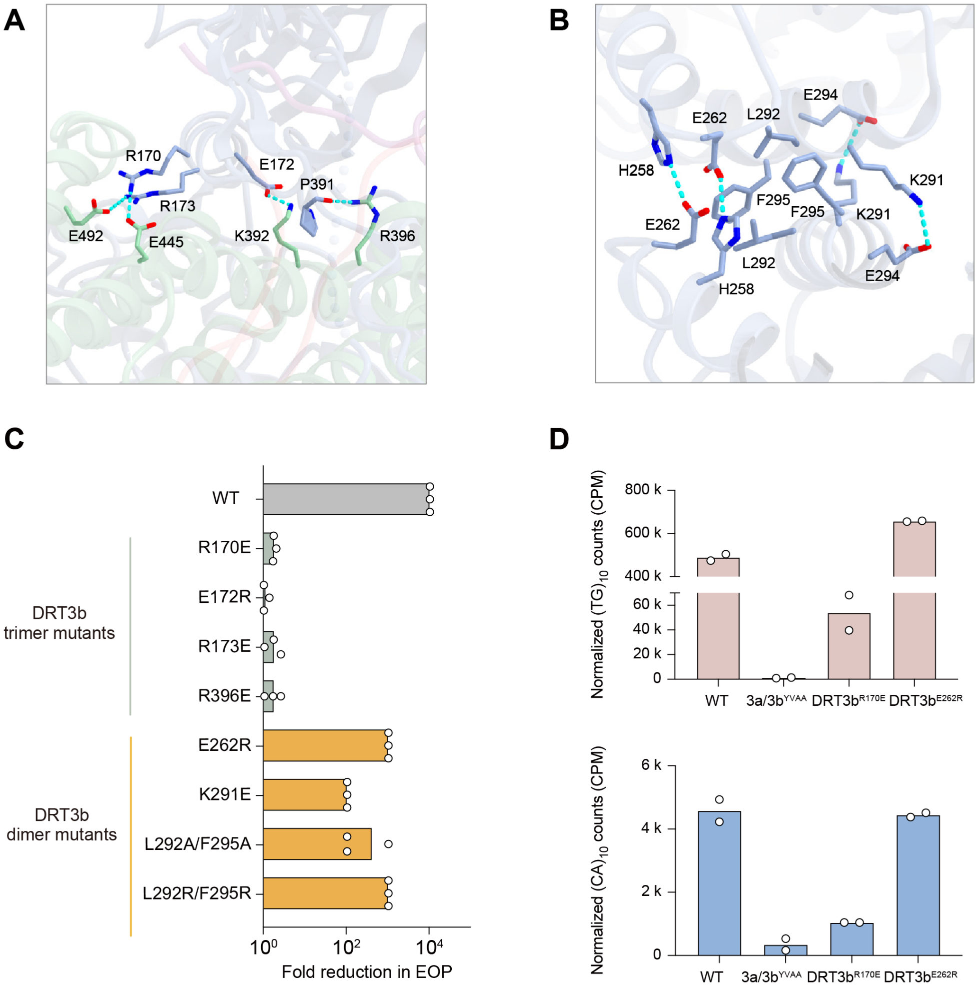


**Figure S7 | DRT3b oligomeric interfaces and their contributions to phage defense and DNA synthesis, Related to Figure 3.**
**(A)** DRT3b trimer interface, highlighting residues that mediate inter-protomer contacts between the αRep domain of one subunit and the RT-like domain of a neighboring subunit. Interacting side chains are shown as sticks and predicted hydrogen-bonding interactions are indicated.
**(B)** DRT3b dimer-of-trimers interface, highlighting residues that contribute to contacts between the two trimers within the DRT3b hexamer. Interacting side chains and predicted hydrogen-bonding interactions are shown as in **A**.
**(C)** Functional analysis of DRT3b interface mutants, quantified as the fold reduction in efficiency of plating (EOP) relative to an EV control. Mutations targeting the trimer interface abolish or strongly impair defense, whereas mutations targeting the dimer-of-trimers interface retain substantial activity. Data are from $n = 3$ technical replicates.
**(D)** Quantification of normalized cDIP-seq reads containing $(dTdG)_{10}$ (top) and $(dCdA)_{10}$ (bottom) motifs, plotted as counts per million (CPM), for WT and indicated interface mutants. Disruption of the trimer interface abolishes production of both repeat species, whereas dimer-of-trimers interface mutants retain near-WT levels of cDNA products. Data represent the mean with individual data points from $n = 2$ independent biological replicates.

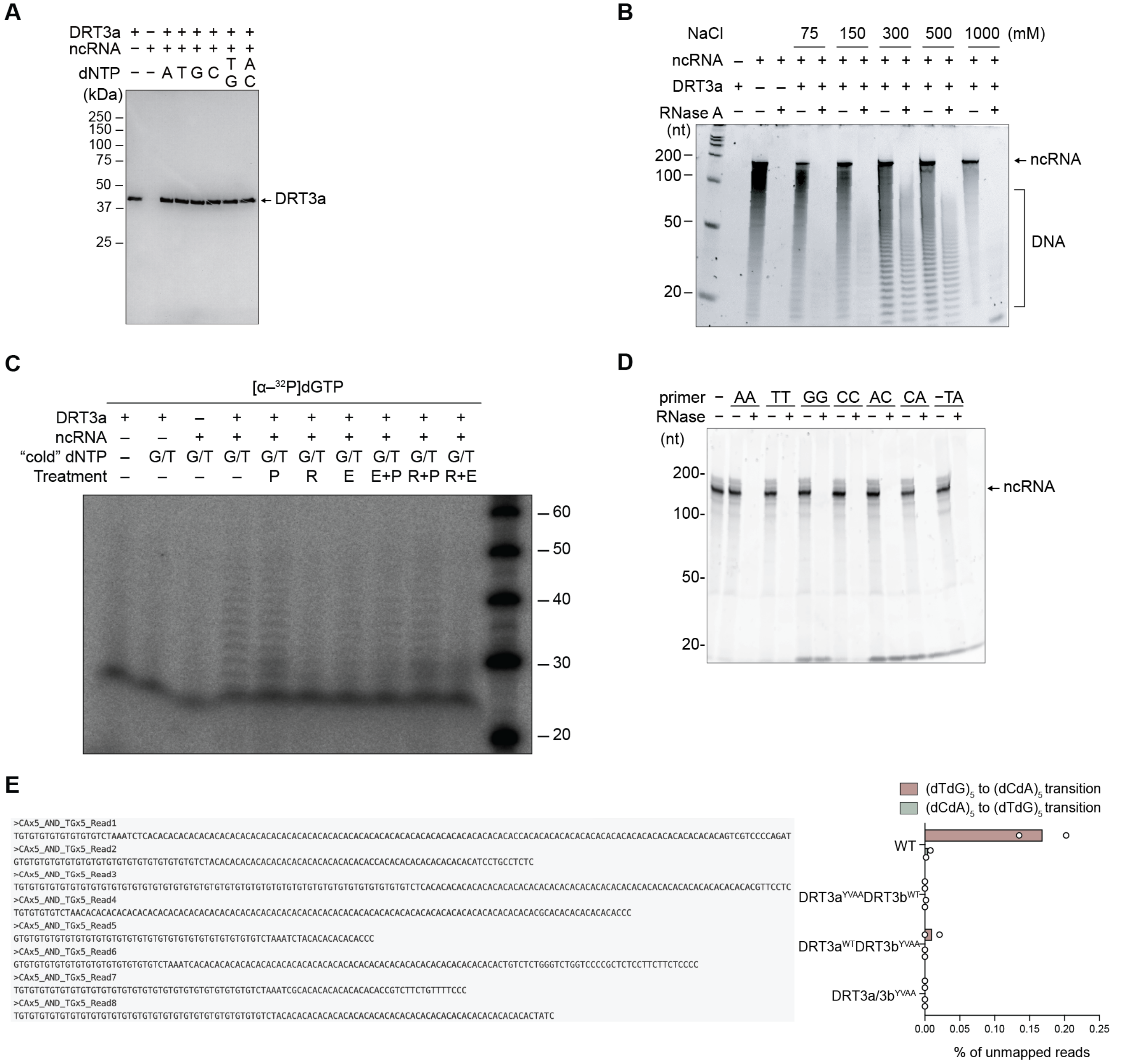


**Figure S8 | DRT3a does not exhibit protein- or primer-dependent initiation, Related to Figure 4.**
**(A)** SDS–PAGE analysis of DRT3a+ncRNA reactions with dGTP/dTTP showing no mobility shift consistent with protein–DNA conjugate formation.
**(B)** Urea–PAGE with SYBR Gold stain analysis showing no evidence of ncRNA shift with production of cDNA.
**(C)** Radiolabeled nucleotide incorporation assays with DRT3a and ncRNA showing no detectable protein priming. Lane labels indicate treatment: P (proteinase K), R (RNase A), E (exonuclease I).
**(D)** Reactions supplemented with 20-nt oligonucleotide primers fail to support extension, indicating a lack of primer-dependent synthesis. Primers included homopolymeric oligos, poly-A (AA), poly-T (TT), poly-G (GG), and poly-C (CC); alternating dinucleotide oligos, poly-AC (AC) and poly-CA (CA); and an oligo complementary to the upstream region of the ncRNA (-TA) (Table S4).
**(E)** *In vitro* reverse transcription sequencing analysis showing absence of CA→GT transitions, consistent with a non–protein-primed, non-templated synthesis mechanism (left). Percentage of unmapped reads that contain either $(dTdG)_5$ to $(dCdA)_5$ transitions or $(dCdA)_5$ to $(dTdG)_5$ transitions (right).

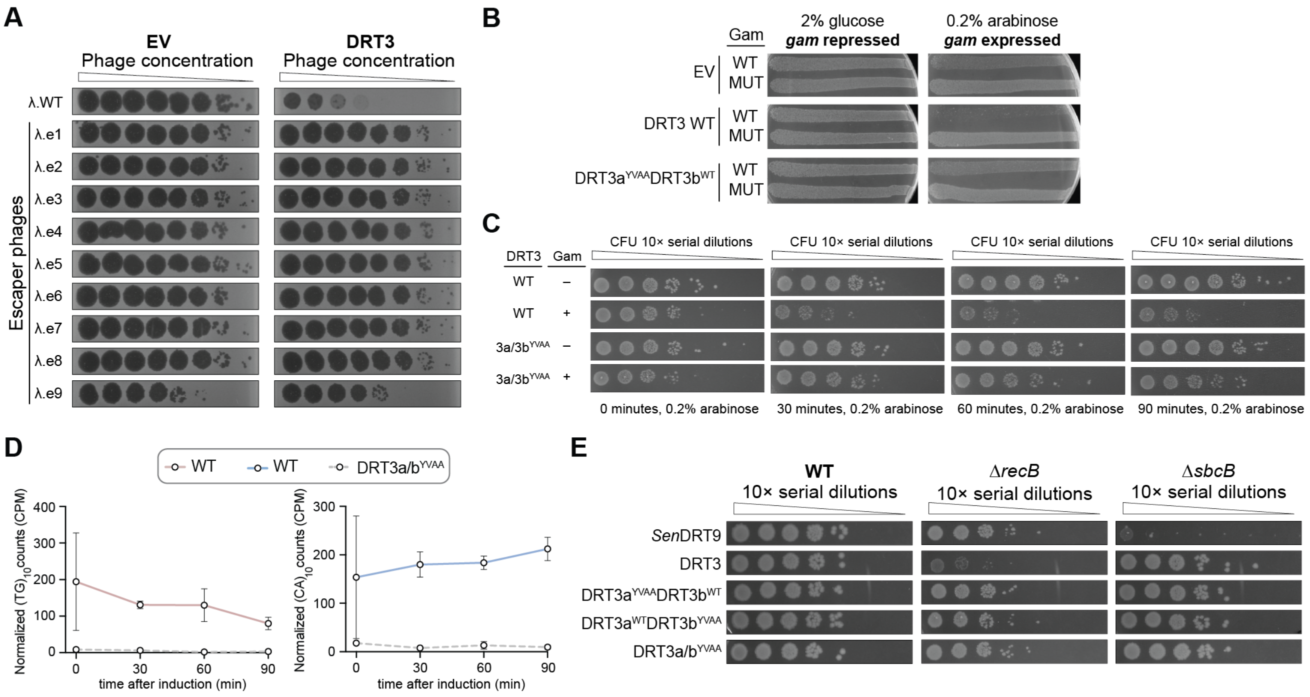


**Figure S9 | Gam-mediated triggering of DRT3 and involvement of RecBCD, Related to Figure 5.**

**(A)** Plaque assays of λ escaper phages on empty vector (EV) control and DRT3-expressing strains, identifying *gam*-dependent escape mutants.

**(B)** Induction of WT Gam, but not mutant Gam, results in cell death in DRT3-expressing cells specifically upon arabinose induction.

**(C)** *E. coli* spot assays for the indicated strains expressing WT or catalytic mutant DRT3, with or without inducible Gam. Serial dilutions reveal that Gam-mediated toxicity requires an intact DRT3 system and is observed only under inducing conditions.

**(D)** Miniprep seq–based quantification shows no detectable changes in poly-(dTdG) and poly-(dCdA) species in *E. coli* strains containing the WT DRT3 system after Gam induction. Data are mean ± SD of $n$ = 3 independent biological replicates.

**(E)** *E. coli* spot assays for the indicated WT, Δ*recB*, or Δ*sbcB* strains transformed with the DRT3 or DRT9 constructs shown on the left. Loss of RecB results in synthetic lethality in the presence of DRT3, whereas disruption of the ssDNA exonuclease ExoI (Δ*sbcB*) has minimal effect, highlighting a specific requirement for RecBCD in regulating DRT3-derived dsDNA products. The opposite relationship holds true for DRT9, which produces ssDNA poly-dA species and requires ExoI for autoimmunity avoidance.

**Figure S10 | Metabolomics, transcriptomics, and functional genomics analyses during DRT3-mediated defense and**

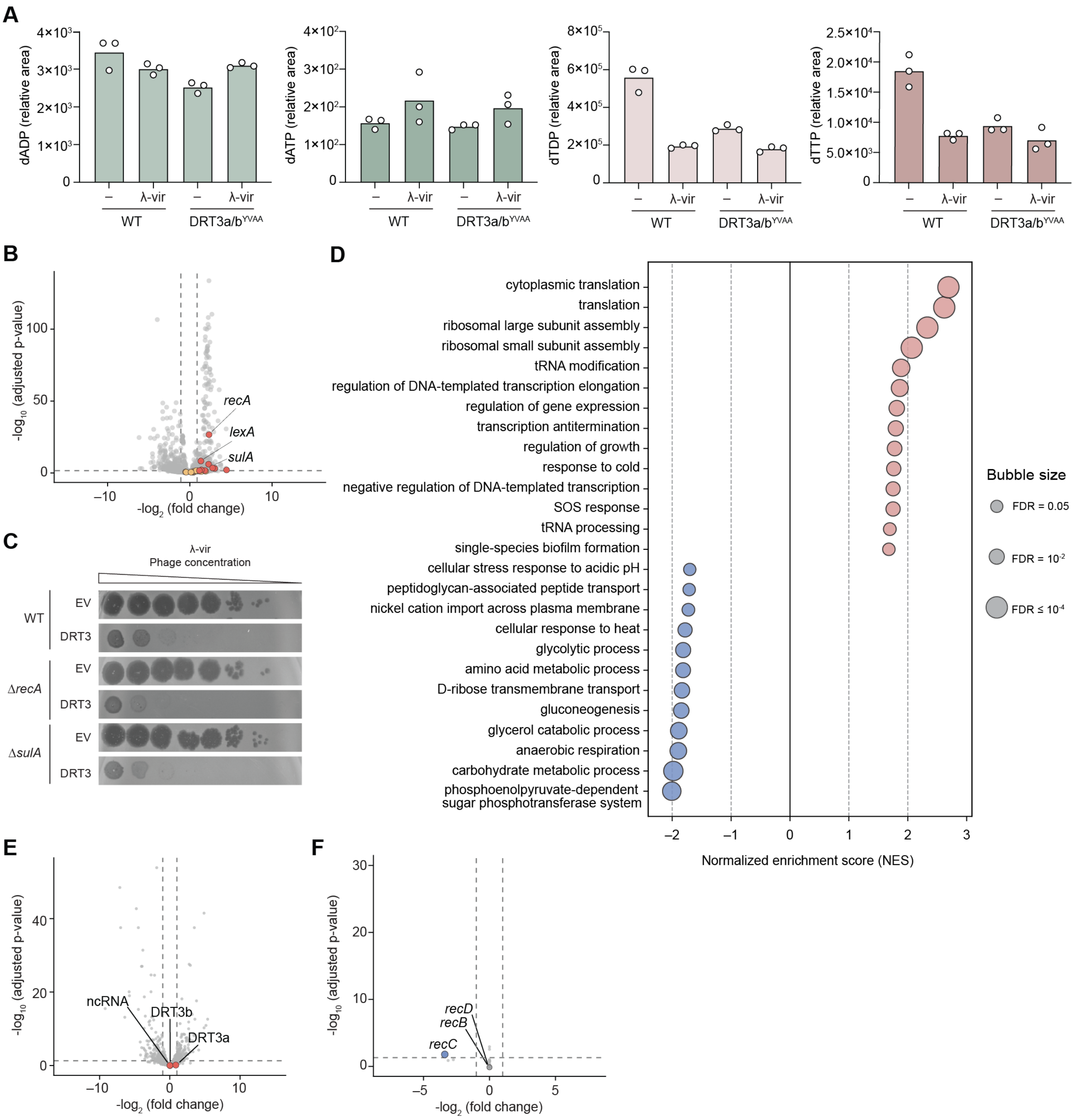


**abortive infection, Related to Figure 5.**

**(A)** Bar graphs of relative nucleotide levels for the indicated species quantified by LC-MS/MS, in lysates from infected and uninfected cells, comparing WT and DRT3a/b$^{YVAA}$ strains.

**(B)** Comparison of transcriptome-wide expression levels between WT and DRT3a/b$^{YVAA}$ strains following λ-vir infection. Colored data points are genes annotated with the GO Biological Process term "SOS response": red points pass both significance and fold-change thresholds, while orange points are SOS-annotated genes that do not. *recA*, *lexA*, and *sulA* transcripts are shown to be enriched in the infected DRT3 WT condition. Dashed lines indicate thresholds for statistical significance ($p_{adj} < 0.05$) and fold-change cutoffs ($|\log_2 FC| > 1$). Data are from $n = 2$ independent biological replicates.

**(C)** Plaque assays of EV and DRT3 in a background Δ*recA* or Δ*sulA* strain do not show loss of defense, suggesting that DRT3-mediated defense is independent of the SOS response pathway.

**(D)** Gene set enrichment analysis (GSEA) of genes ranked by differential expression between WT and DRT3a/b$^{YVAA}$ strains following λ-vir infection. The SOS response gene set was significantly enriched in the WT infected condition (NES = 1.75, $p_{adj} < 0.05$). Data are from *n* = 2 independent biological replicates.

**(E)** Tn–seq $\log_2$ fold-change analysis of DRT3$^{WT}$ + Gam and DRT3a/b$^{YVAA}$ + Gam under glucose Gam suppression highlighting depletion in transposon insertion counts in *recC* and non-significant differential insertion counts in *recB* and *recD*. Dashed lines indicate thresholds for statistical significance ($p_{adj} < 0.05$) and fold-change cutoffs ($|\log_2 FC| > 1$). Data are from *n* = 5 independent biological replicates.

**(F)** Comparison of DRT3a, DRT3b, and ncRNA transcript levels between *E. coli* WT and Δ*greA* strains shows no differential enrichment. Dashed lines indicate thresholds for statistical significance ($p_{adj} < 0.05$) and fold-change cutoffs ($|\log_2 FC| > 1$). Data are from *n* = 3 independent biological replicates.

**Figure S11 | Predicted structural similarity between phage T1 protein ST61 and cellular transcription elongation factors,**

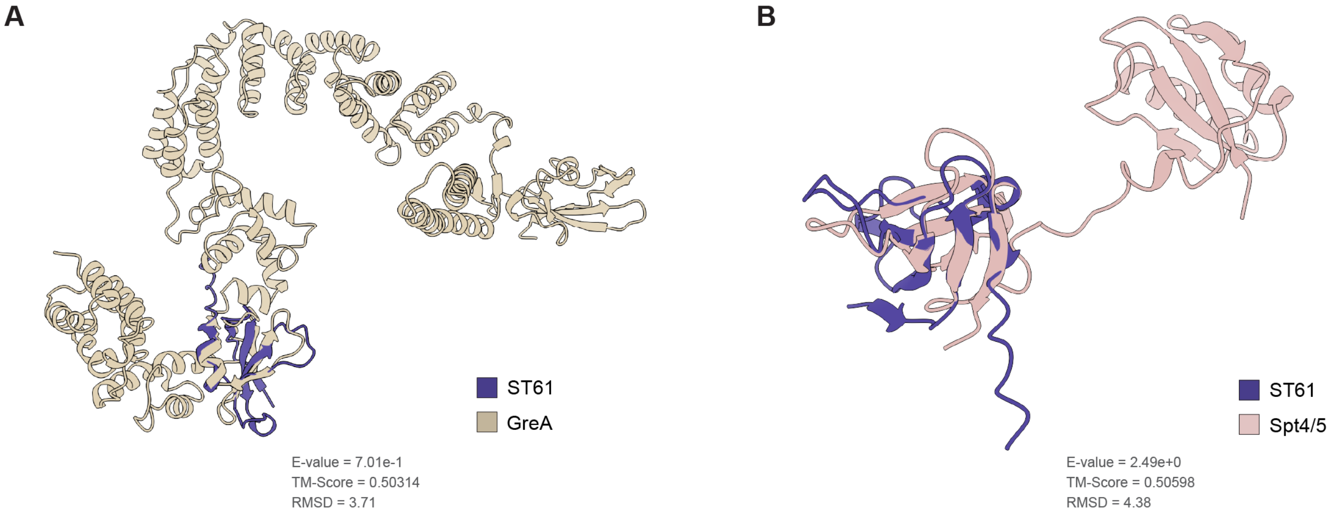


**Related to Figure 5.**

**(A)** Structural alignment of ST61 AlphaFold3 prediction with transcription elongation factor GreA from *Chlamydia muridarum* str. Nigg, generated by FoldSeek (AFDB-Swiss-Prot database).

**(B)** Structural alignment of ST61 AlphaFold3 prediction with transcription elongation factor Spt4/5 from *Pyrococcus furiosus* (PDB ID: 3P8B), generated by FoldSeek.

#### Table S2 | Strains used in this study, Related to STAR Methods.

| Strain ID | Description | NCBI accession (and/or description) | Source |
|---|---|---|---|
| sSL0810 | E. coli MG1655 | U00096.3 | Yale *E. coli* Genetic Stock Center |
| sSL0410 | E. coli NEB Turbo | Escherichia coli strain from New England Biolabs (catalogue #C2984H) | New England BioLabs (C2984H) |
| sSL0800 | E. coli BW25113 | $F^-$, Δ(*araD-araB*)567, Δ*lacZ*4787(::*rrnB*-3), $\lambda^-$, *rph*-1, Δ(*rhaD-rhaB*)568, *hsdR*514 | Gift of S. Tavazoie from Horizon Discovery Keio Collection |
| sSL5293 | E. coli BW25113 Δ*sbcB* | $F^-$, Δ(*araD-araB*)567, Δ*lacZ*4787(::*rrnB*-3), $\lambda^-$, Δ*sbcB*780::*kan*, *rph*-1, Δ(*rhaD-rhaB*)568, *hsdR*514 | Gift of S. Tavazoie from Horizon Discovery Keio Collection |
| sSL6348 | E. coli BW25113 Δ*recA* | $F^-$, Δ(*araD-araB*)567, Δ*lacZ*4787(::*rrnB*-3), $\lambda^-$, Δ*recA*774::*kan*, *rph*-1, Δ(*rhaD-rhaB*)568, *hsdR*514 | Horizon Discovery Keio Collection |
| sSL0806 | E. coli BW25113 Δ*recB* | $F^-$, Δ(*araD-araB*)567, Δ*lacZ*4787(::*rrnB*-3), $\lambda^-$, Δ*recB*745::*kan*, *rph*-1, Δ(*rhaD-rhaB*)568, *hsdR*514 | Gift of S. Tavazoie from Horizon Discovery Keio Collection |
| sSL0807 | E. coli BW25113 Δ*recC* | $F^-$, Δ(*araD-araB*)567, Δ*lacZ*4787(::*rrnB*-3), $\lambda^-$, Δ*recC*747::*kan*, *rph*-1, Δ(*rhaD-rhaB*)568, *hsdR*514 | Gift of S. Tavazoie from Horizon Discovery Keio Collection |
| sSL0805 | E. coli BW25113 Δ*recD* | $F^-$, Δ(*araD-araB*)567, Δ*lacZ*4787(::*rrnB*-3), $\lambda^-$, Δ*recD*744::*kan*, *rph*-1, Δ(*rhaD-rhaB*)568, *hsdR*514 | Gift of S. Tavazoie from Horizon Discovery Keio Collection |
| sSL6352 | E. coli BW25113 Δ*sulA* | $F^-$, Δ(*araD-araB*)567, Δ*lacZ*4787(::*rrnB*-3), $\lambda^-$, Δ*sulA*773::*kan*, *rph*-1, Δ(*rhaD-rhaB*)568, *hsdR*514 | Horizon Discovery Keio Collection |
| sSL6000 | E. coli BW25113 Δ*greA* | $F^-$, Δ(*araD-araB*)567, Δ*lacZ*4787(::*rrnB*-3), $\lambda^-$, Δ*greA*788::*kan*, *rph*-1, Δ(*rhaD-rhaB*)568, *hsdR*514 | Horizon Discovery Keio Collection |
| sHN0002 | E. coli Rosetta2 (DE3) | $F^-$, *ompT*, *hsdS*$_B$ ($r_B^-$ $m_B^-$), *gal*, *dcm*, (DE3), and pRARE2 ($Cam^R$) | Novagen, USA |

#### Table S4 | Oligonucleotides used in this study, Related to STAR Methods.

| Oligo ID | Description | Sequence (5' to 3') |
|---|---|---|
| oSL21872 | Spike-in oligo for sequencing | TGCTTTGGCTCCGAGCGCTTGGTATTCATGCGCCTATCTTGTG GCCCAAGTCCGAGCTCACATGCCATGCTTTCCCTGCTTGGAT GGGACCTTGCAGCCAATGGCACAACACCATGTGGTTGCTAT AGTTGCAAAAGAGTTGCACCTAAA |
| oSL21873 | Spike-in oligo for sequencing | TAAGGAATTCGGGTACATGAGTCGCTCAATCCGTGCCTTTGT CATTACGGTTTGGTCCTGCCTGTCTCCGTTGCGATGCGTTGTC GGCACATTGTCTATAAAGAACGACTTACTCGAGAGGCTCCA AACAACTACTCGACCTTTAACCGC |
| oSL21874 | Spike-in oligo for sequencing | TGTGCCGTAACTGTCTCGCCTAGCTATACTGCCTTGGTATATC GGCGGCAATTTTAATATATCTCTACACTCTGGACAAGCGTGA AAAGATATAGTCATGCACACTGGGCCTACATCTTGGCTCAAT GGACGTCTCGTTGACAATCGTCC |
| oSL12903 | Top strand R1 oligo for pre-loading TnY | TCGTCGGCAGCGTCAGATGTGTATAAGAGACAG |
| oSL12904 | Top strand R2 oligo for pre-loading TnY | GTCTCGTGGGCTCGGAGATGTGTATAAGAGACAG |
| oSL11032 | Bottom strand oligo for pre-loading TnY | CTGTCTCTTATACA-ddC |
| Nextera index 1 | Oligo with a barcode to amplify tagmentation products | AATGATACGGCGACCACCGAGATCTACAC NNNNNNNN TCGTCGGCAGCGTC |
| Nextera index 2 | Oligo with a barcode to amplify tagmentation products | CAAGCAGAAGACGGCATACGAGAT NNNNNNNN GTCTCGTGGGCTCGG |
| oKYA01 | Template oligo for *in vitro* transcription | CGGGGATCGCAGTGGTGAGTAACCATGCATCATCAGGAGTA CGGATAAAATGCTTGATGGTCGGAAGAGGCATAAATTCCGT CAGCCAGTTTAGTCTGACCATCTCATCTGTAACGGATCCTAA TACGACTCACTATAGGTTAAAGGGTTGCGATGCCTAAGGTTT CGACCTGAAGCAGATACCGGAAGATCGGCTTTTGAATGTTC ATCCGAAAGATATTCGCGATACGTTTTGAGGATGGACCGATT TAGACACACTATTGCCTTTTAGCTAAACAGGCCGCGAAAGC GGCCTTTTTA |
| oKYA02 | poly-A oligo | AAAAAAAAAAAAAAAAAAAA |
| oKYA03 | poly-T oligo | TTTTTTTTTTTTTTTTTTTT |
| oKYA04 | poly-G oligo | GGGGGGGGGGGGGGGGGGGG |
| oKYA05 | poly-C oligo | CCCCCCCCCCCCCCCCCCCC |
| oKYA06 | poly-AC oligo | ACACACACACACACACACAC |
| oKYA07 | poly-CA oligo | CACACACACACACACACACA |
| oKYA08 | Oligo containing a sequence complementary to the upstre | CACACACACACACACCAATA |

#### Table S5 | Cryo-EM data collection, refinement, and validation statistics, Related to Figure 3.

| DRT3b hexamer \| EMDB EMD-80550 \| PDB 26CZ | |
|---|---|
| **Data collection and processing** | |
| Microscope | Titan Krios G3i |
| Detector | K3 camera |
| Automation software | EPU |
| Magnification | 105000 |
| Voltage (kV) | 300 |
| Energy filter slit width (eV) | 2 |
| Exposure rate ($e^{-}$/Å$^{2}$/s) | 10.9 |
| Total electron exposure ($e^{-}$/Å$^{2}$) | 50.6 |
| Defocus range (μm) | −0.8 to −2.0 |
| Pixel size (Å) | 0.83 |
| Number of frames per image | 50 |
| Number of micrographs | 835 |
| Symmetry imposed | D3 |
| Initial particle images (no.) | 35460 |
| Final particle images (no.) | 13084 |
| Map resolution (Å) | 3.1 |
| FSC threshold | 0.143 |
| **Model building and refinement** | |
| Model composition | |
| (Model composition in the asymmetric unit) | |
| peptide residues | 640 |
| RNA residues | 0 |
| DNA residues | 14 |
| Protein atoms | 5,299 |
| DNA atoms | 280 |
| Other atoms | 7 |
| Average B factors (Å$^{2}$) | |
| peptide B-factors | 105.1 |
| DNA B-factors | 124.4 |
| other B-factors | 66.4 |
| R.M.S. deviations from ideal | |
| bond lengths rmsd (Å) | 0.0065 |
| bond angles rmsd (°) | 1.12 |
| Validation | |
| Clashscore | 0.6 |
| Rotamer outliers (%) | 3.03 |
| Ramachandran plot (%) | |
| Ramachandran favored | 93.99 |
| Ramachandran allowed | 5.7 |
| Ramachandran outliers | 0.32 |

#### Table S6 | Genotypes of escaper phages that evade *Eco*DRT3 immunity, Related to Figure 5.

| Escaper ID | Phage | Mutated/lost genes | Mutation start coordinate in phage genome | Mutation | Predicted effect of mutation |
|---|---|---|---|---|---|
| λ.e1 | λ | *gam* -like host nuclease inhibitor | 26,879 | coding (359/417 nt) | +A |
| λ.e2 | λ | *gam* -like host nuclease inhibitor | 26,952 | coding (286/417 nt) | +C |
| λ.e3 | λ | central tail fiber *J* | 18,623 | M1040T (ATG→ACG) | T→C |
| | | central tail fiber *J* | 18,823 | D1107N (GAT→AAT) | G→A |
| | | *gam* -like host nuclease inhibitor | 26,943 | coding (295/417 nt) | (T)6→7 |
| λ.e4 | λ | *gam* -like host nuclease inhibitor | 27,050 | coding (188/417 nt) | +G |
| λ.e5 | λ | *gam* -like host nuclease inhibitor | 27,081 | Q53* (CAG→TAG) | G→A |
| λ.e6 | λ | *gam* -like host nuclease inhibitor | 26,943 | coding (295/417 nt) | (T)6→7 |
| λ.e7 | λ | *gam* -like host nuclease inhibitor | 26,997 | Q81* (CAG→TAG) | G→A |
| λ.e8 | λ | *gam* -like host nuclease inhibitor | 27,011 | coding (227/417 nt) | (T)5→6 |
| λ.e9 | λ | *gam* -like host nuclease inhibitor | 26,879 | coding (359/417 nt) | +A |
