## Supplementary material for "Coordinated RNA- and protein-templated synthesis of double-stranded DNA by a dual reverse transcriptase immune system": Key Resources Table

| REAGENT or RESOURCE | SOURCE | IDENTIFIER |
|---|---|---|
| **Antibodies** | | |
| Mouse monoclonal anti-FLAG | Sigma-Aldrich | Cat#F3165; RRID: AB_259529 |
| **Bacterial and virus strains** | | |
| *E. coli* K-12 MG1655 (sSL0810) | Yale *E. coli* Genetic Stock Center | U00096.3 |
| *E. coli* NEB Turbo (sSL0410) | New England Biolabs | Cat#C2984H |
| *E. coli* BW25113 (sSL0800) | Gift of S. Tavazoie from Horizon Discovery Keio Collection[40] (Baba et al., 2006) | CGSC#7636 |
| *E. coli* BW25113 Δ*sbcB* (sSL5293) | Gift of S. Tavazoie from Horizon Discovery Keio Collection[40] (Baba et al., 2006) | JW1993 |
| *E. coli* BW25113 Δ*recA* (sSL6348) | Horizon Discovery Keio Collection[40] (Baba et al., 2006) | JW2669 |
| *E. coli* BW25113 Δ*recB* (sSL0806) | Gift of S. Tavazoie from Horizon Discovery Keio Collection[40] (Baba et al., 2006) | JW2788 |
| *E. coli* BW25113 Δ*recC* (sSL0807) | Gift of S. Tavazoie from Horizon Discovery Keio Collection[40] (Baba et al., 2006) | JW2790 |
| *E. coli* BW25113 Δ*recD* (sSL0805) | Gift of S. Tavazoie from Horizon Discovery Keio Collection[40] (Baba et al., 2006) | JW2787 |
| *E. coli* BW25113 Δ*sulA* (sSL6352) | Horizon Discovery Keio Collection[40] (Baba et al., 2006) | JW0941 |
| *E. coli* BW25113 Δ*greA* (sSL6000) | Horizon Discovery Keio Collection[40] (Baba et al., 2006) | JW3148 |
| Bacteriophage λ-vir | Gift from M. Laub | N/A |
| Bacteriophage T5 | Gift from M. Laub | N/A |
| *E. coli* Rosetta2 (DE3) (sHN0002) | Novagen | Cat#71397 |
| **Chemicals, peptides, and recombinant proteins** | | |
| DRT3a protein | This paper | N/A |
| DRT3a mutant protein | This paper | N/A |
| DRT3b protein | This paper | N/A |
| DRT3b mutant proteins | This paper | N/A |
| SUMO protease | This paper | N/A |
| T7 RNA polymerase | This paper | N/A |
| Proteinase K for *in vitro* gel experiments | Sigma-Aldrich | Cat#3115836001 |
| RNase A for *in vitro* gel experiments | NEB | Cat#T3018L |
| Proteinase K for *in vivo* and *in vitro* sequencing experiments | Thermo Fisher Scientific | Cat#EO0491 |
| RNase A for *in vivo* and *in vitro* sequencing experiments | Thermo Fisher Scientific | Cat#EN0531 |
| TnY transposase | This paper | N/A |
| TURBO DNase | Thermo Fisher Scientific | Cat#AM2238 |
| RppH | New England Biolabs | Cat#M0356 |
| T4 polynucleotide kinase | New England Biolabs | Cat#M0201 |
| DNase I | New England Biolabs | Cat#M0303 |
| Exonuclease I | New England Biolabs | Cat#M0568 |
| MmeI | New England Biolabs | Cat#R0637 |
| Sequencing-grade modified trypsin | Promega | Cat#V5113 |
| SUPERase•In RNase Inhibitor | Thermo Fisher Scientific | Cat#AM2696 |
| **Critical commercial assays** | | |
| xGen ssDNA & Low-Input DNA Library Prep Kit | Integrated DNA Technologies | Cat#10009817 |
| HyperLight FluorGreen dsDNA Assay Kit | APExBIO | Cat#K1603 |
| **Deposited data** | | |
| Hexameric DRT3b complex map | This paper | EMD-80550 |
| Hexameric DRT3b complex coordinates | This paper | PDB: 26CZ |
| Raw cryo-EM images of the hexameric DRT3b complex | This paper | EMPIAR-13689 |
| λ escaper sequencing | This paper | BioProject: PRJNA1461384 |
| cDIP–sequencing | This paper | GEO: GSE330639 |
| Miniprep–sequencing | This paper | GEO: GSE330641 |
| Tn–sequencing | This paper | GEO: GSE329894 |
| RNA–sequencing | This paper | GEO: GSE329895 |
| RIP–sequencing | This paper | GEO: GSE329897 |
| Immunoprecipitation-mass spectrometry | This paper | MassIVE: MSV000101674 |
| **Oligonucleotides** | | |
| DNA oligos (for *in vitro* transcription sequencing) | This paper | Table S4 |
| DNA oligos (for TnY escaper sequencing) | This paper | Table S4 |
| DNA oligos (for *in vitro* transcription) | This paper | Table S4 |
| DNA oligos (for *in vitro* assays) | This paper | Table S4 |
| **Recombinant DNA** | | |
| pSL0007-pSL11244 | This paper | Table S1 |
| pKYA01-pKYA11 | This paper | Table S1 |
| **Software and algorithms** | | |
| EPU software | Thermo Fisher Scientific | https://www.thermofisher.com/jp/en/home/electron-microscopy/products/software-em-3d-vis/epu-software.html |
| cryoSPARC v5.0.2 | Punjani et al.[60] | https://cryosparc.com/; RRID:SCR_016501 |
| COOT | Emsley et al.[65] | https://www2.mrc-lmb.cam.ac.uk/personal/pemsley/coot/; RRID:SCR_014222 |
| Boltz-2 | Passaro et al.[66] | https://github.com/jwohlwend/boltz |
| Servalcat | Yamashita et al.[67] | https://github.com/keitaroyam/servalcat |
| ProSMART | Nicholls et al.[68] | https://www.ccp4.ac.uk/html/prosmart.html |
| MolProbity | Williams et al.[69] | https://www.phenix-online.org/documentation/reference/molprobity_tool.html; RRID:SCR_014226 |
| Topaz | Bepler et al.[61] | https://github.com/tbepler/topaz |
| UCSF ChimeraX | Pettersen et al.[70] | https://www.rbvi.ucsf.edu/chimerax/; RRID:SCR_015872 |
| CueMol | N/A | http://www.cuemol.org |
| Prism | Graphpad | https://www.graphpad.com/scientific-software/prism/ |

| REAGENT or RESOURCE | SOURCE | IDENTIFIER |
|---|---|---|
| **Software and algorithms** | | |
| Cutadapt v5.0 | Martin[50] | https://cutadapt.readthedocs.io; RRID:SCR_011841 |
| bwa-mem2 v2.2.1 | Vasimuddin et al.[51] | https://github.com/bwa-mem2/bwa-mem2; RRID:SCR_022192 |
| Bowtie2 v2.2.1 | Langmead and Salzberg[73] | http://bowtie-bio.sourceforge.net/bowtie2; RRID:SCR_016368 |
| SAMtools v1.17 | Danecek et al.[52] | http://www.htslib.org; RRID:SCR_002105 |
| deepTools (bamCoverage) v3.5.6 | Ramírez et al.[53] | https://deeptools.readthedocs.io; RRID:SCR_016366 |
| featureCounts (Subread) v2.0.2 | Liao et al.[55] | https://subread.sourceforge.net; RRID:SCR_012919 |
| PyDESeq2 v0.5.3 | Muzellec et al.[56] | https://github.com/owkin/PyDESeq2 |
| gseapy v1.1.13 | Fang et al.[77] | https://github.com/zqfang/GSEApy |
| bbduk (BBTools) v39.83 | Bushnell[76] | https://sourceforge.net/projects/bbmap/ |
| breseq v0.39.0 | Deatherage and Barrick[75] | https://github.com/barricklab/breseq; RRID:SCR_010810 |
| MEME v5.5.7 | Bailey et al.[58] | https://meme-suite.org; RRID:SCR_001783 |
| IGV v2.17.4 | Robinson et al.[54] | https://igv.org; RRID:SCR_011793 |
| MaxQuant v2.0.3.0 | Cox and Mann[59] | https://www.maxquant.org; RRID:SCR_014485 |
| Skyline-daily v24.1.1.398 | MacLean et al.[79] | https://skyline.ms; RRID:SCR_014080 |
| MMseqs2 | Steinegger and Söding[41] | https://github.com/soedinglab/MMseqs2 |
| MAFFT v7.490 | Katoh and Standley[45] | https://mafft.cbrc.jp/alignment/software/; RRID:SCR_011811 |
| FastTree 2 v2.1.11 | Price et al.[43] | http://www.microbesonline.org/fasttree/; RRID:SCR_015501 |
| iTOL | Letunic and Bork[44] | https://itol.embl.de; RRID:SCR_018174 |
| mLocARNA v2.0.1 | Will et al.[46] | http://www.bioinf.uni-freiburg.de/Software/LocARNA/ |
| Infernal v1.1.4 | Nawrocki and Eddy[47] | http://eddylab.org/infernal/; RRID:SCR_011809 |
| CD-HIT | Fu et al.[48] | https://sites.google.com/view/cd-hit; RRID:SCR_007105 |
| R-scape v2.6.10 | Rivas et al.[49] | http://eddylab.org/R-scape/ |
| DECIPHER v3.8.0 | Wright[71] | https://bioconductor.org/packages/DECIPHER/; RRID:SCR_001357 |
| WebLogo v3.7.9 | Crooks et al.[72] | https://weblogo.threeplusone.com; RRID:SCR_010236 |
| Xcalibur v4.5.474.0 | Thermo Fisher Scientific | https://www.thermofisher.com/us/en/home/industrial/mass-spectrometry/liquid-chromatography-mass-spectrometry-lc-ms/lc-ms-software/lc-ms-data-acquisition-software/xcalibur-data-acquisition-interpretation-software.html; RRID:SCR_014593 |
| **Other** | | |
| Ni-NTA Superflow resin | QIAGEN | Cat#30450 |
| HiTrap Heparin HP column | GE Healthcare | Cat#17040601 |
| Superdex 200 Increase 10/300 column | GE Healthcare | Cat#28990944 |
| Amicon Ultra-4 mL Centrifugal Filter Unit (MWCO 30 kDa) | Merck Millipore | Cat#UFC803008 |
| Amicon Ultra-0.5 mL Centrifugal Filter (MWCO 3 kDa) | Merck Millipore | Cat#UFC5003 |
| Cu 300 mesh R1.2/1.3 grid covered with a 2 nm carbon film | Quantifoil | https://www.quantifoil.com/products/quantifoil/quantifoil-circular-holes/ |
| Protein G Dynabeads | Thermo Fisher Scientific | Cat#10004D |
